# Optimization of methods for rapid and robust generation of cardiomyocyte-specific *crispants* in zebrafish using the *cardiodeleter* system

**DOI:** 10.1101/2024.09.27.615502

**Authors:** Sean Keeley, Miriam Fernández-Lajarín, David Bergemann, Nicolette John, Lily Parrott, Brittany E. Andrea, Juan Manuel González-Rosa

## Abstract

CRISPR/Cas9 has massively accelerated the generation of gene loss-of-function models in zebrafish. However, establishing tissue-specific mutant lines remains a laborious and time-consuming process. Although a few dozen tissue-specific Cas9 zebrafish lines have been developed, the lack of standardization of some key methods, including gRNA delivery, has limited the implementation of these approaches in the zebrafish community. To tackle these limitations, we have established a cardiomyocyte-specific Cas9 line, the *cardiodeleter*, which efficiently generates biallelic mutations in combination with gene-specific gRNAs. We have also optimized the development of transposon-based *guide shuttles* that carry gRNAs targeting a gene of interest and permanently label the cells susceptible to becoming mutant. We validated this modular approach by deleting five genes (*ect2*, *tnnt2a*, *cmlc2*, *amhc*, and *erbb2*), all resulting in the loss of the corresponding protein or phenocopying established mutants. Additionally, we provide detailed protocols describing how to generate *guide shuttles*, which will facilitate the dissemination of these techniques in the zebrafish community. Our approach enables the rapid generation of tissue-specific *crispants* and analysis of mosaic phenotypes, bypassing limitations such as embryonic lethality, making it a valuable tool for cell-autonomous studies and genetic screenings.

## INTRODUCTION

In the last three decades, zebrafish have been exploited as a powerful model system to identify the genes and mechanisms that orchestrate cardiovascular development in vertebrates^1^. The pioneering forward screening of cardiovascular phenotypes in the 1990s uncovered hundreds of genes involved in cardiac development and heart function^2,3^. Implementing reverse genetic strategies, including zinc-finger nucleases^4^, TALENs^5^, and most recently, CRISPR/Cas^6^, has re-invigorated the field. The ability to mutate specific genes is critical nowadays when massive datasets are generated yearly from single-cell sequencing and genome-wide association studies (GWAS). To move from correlations (i.e., spatial expression or association between mutations and specific pathologies), we need experimental systems that enable probing gene function *in vivo* with moderate or high throughput.

The generation of zebrafish mutants by injection of Cas9 and guide RNAs (gRNAs) targeting a gene of interest (GOI) has become an invaluable reverse genetic approach^7^. The efficiency of mutagenesis using this methodology is very high, and often the F_0_-injected animals, called *crispants*^8^, exhibit the mutant phenotype. This strategy allows researchers to determine loss-of-function (LOF) phenotypes in just a few days, eliminating the laborious process of establishing a mutant strain. In the last few years, *crispant* screens have become very popular^9–11^, even though they present two important and intertwined limitations. First, *crispants* are mosaics composed of wild-type and mutant cells, but there is no labeling that would distinguish between these populations, and the percentage of mutant cells is highly variable between animals. Second, because the mutations are not restricted to a specific cell type, it is almost impossible to attribute a given phenotype to the loss of the GOI in a specific tissue or cell population. For example, an elegant study recently reported the development of cavernous vascular malformations in *ccm2 crispants*, but it remains unknown whether this phenotype emerges from the loss of this gene in the vascular endothelium or other cell types^12^. Undoubtedly, the gold standard to dissect the role of a gene in a given tissue involves the use of the Cre/lox technology, and while recent efforts have improved the efficiency of the generation of these conditional alleles in zebrafish^13,14^, these methods are costly and time-consuming, severely reducing throughput.

Tissue-specific Cas9 transgenic lines have been developed as an alternative to induce LOF mutations in defined cell populations^15^. In the context of the heart, transgenic mice expressing Cas9 exclusively in the myocardium have been established^16,17^ and are now regularly used to induce cardiomyocyte-specific LOF models rapidly and efficiently^18–20^. A myocardial-Cas9 transgene is stably inserted in the genome in these animals, and the gRNAs are delivered via adeno-associated virus 9 (AAV9) infection. By separating Cas9 and the gRNAs, this system becomes modular: the same stable Cas9 transgenic strain can be treated with AAV9s targeting different GOIs. Moreover, AAV9-infected cardiomyocytes expressing the gRNAs also produce a fluorescent protein, enabling direct identification of the mutant cells. While tissue-specific Cas9 transgenes were adapted in zebrafish almost a decade ago^21,22^, a system with the characteristics described above (i.e., modularity and permanent labeling of mutant cells) has yet to be developed. This is partly due to the lack of viral approaches to deliver the gRNAs to zebrafish. Notably, the implementation of tissue-specific methods in zebrafish has been relatively modest and mainly limited to cancer research.

In contrast to the modular system described in mice, the original method to induce tissue-specific mutations in zebrafish combined Cas9 and a single U6-gRNA cassette in a Tol2-based transgene^21^. This construct included a transgenesis reporter to identify animals carrying the transgene but not specifically labeling cells susceptible to being edited. By design, this approach requires assembling a new Cas9-U6-gRNA construct for each GOI. Moreover, given that the Cas9 coding sequence spans over ∼4 kb, these transgenes tend to be large, reducing the transgenesis efficiency. Although several variations on this method have been reported to date, including attempts to separate Cas9 and the gRNAs in two different transgenes, most strategies require the identification of stable lines expressing each construct^22^. As an alternative to current methods, we present here new genetic tools and protocols to generate cardiomyocyte-specific *crispants*. We leveraged an orthogonal transposase approach to mobilize transposons encoding gRNAs, which we call *guide shuttles*, without disrupting the tissue-specific Cas9 transgene, bypassing an outstanding technical limitation. Because our *guide shuttles* also encode a fluorescent protein, this method labels presumptively mutant cells. We provide an extensive validation of the system by deleting five genes (*ect2*, *tnnt2a*, *cmlc2*, *amhc*, and *erbb2*), all resulting in the loss of the corresponding protein or phenocopying established mutants. We provide a side-by-side comparison of the mutagenesis efficiency using a different number of guides. Additionally, we created detailed protocols to generate *guide shuttles*, which will facilitate the implementation of tissue-specific Cas9 approaches in the zebrafish community.

### Design

The zebrafish community currently lacks a standardized modular approach that allows for generating tissue-specific *crispants* that (1) circumvent the time-consuming process of identifying stable lines and (2) label the population of presumptively edited cells. These limitations prompted us to establish a standardized protocol for a modular, tissue-specific gene disruption system that would allow the permanent labeling of presumptively mutant clones. A system that fulfills these characteristics would be invaluable in determining the cell-autonomous roles of a gene, replacing the use of blastomere transplantations^23–25^. We reasoned that a cardiomyocyte-specific Cas9 line could serve as the springboard for validating and optimizing the conditions to generate tissue-specific *crispants*. The zebrafish heart is accessible and can be easily imaged in the embryo, and decades of research have identified genes that, when mutated, result in specific phenotypes that could be used to validate this approach. Separating the Cas9 and gRNA components would allow us to implement an intersectional, double-labeling system using different fluorescent proteins. By inspecting embryos under a fluorescent scope, a researcher can identify animals that carry Cas9 and have clones expressing the guides, discarding unsuccessful injections. The LOF phenotypes may be evaluated a few days post-injection, transforming this approach into a viable screening strategy. Moreover, because of the mosaic nature of the gene disruption, clones of mutant cardiomyocytes may survive to adulthood intermingled with wild-type cells, bypassing the embryonic lethality that would occur in cases where the gene is absent in the entire embryo. Promising candidates that elicit the desired phenotype can be raised and screened for transmission, and then a stable line carrying the gRNAs can be identified for additional studies.

## RESULTS

### The *cardiodeleter* transgene efficiently induces cardiomyocyte-specific mutations in targeted genes

To induce cardiomyocyte-specific CRISPR/Cas9-mediated mutations, we designed a transgene in which the cardiomyocyte-specific promoter *cmlc2* drives the expression of a bicistronic cassette encoding nuclear GFP and a zebrafish codon-optimized, nuclear-directed form of Cas9 **(Figure 1A)**. This Cas9 version has been used to efficiently induce double-strand breaks by mRNA injection into fertilized zebrafish oocytes^26^. Animals carrying this transgene, called the *cardiodeleter*, developed normally to adulthood and exhibited morphologically normal hearts with strong expression of GFP and Cas9 in cardiomyocytes **(Figure 1B and Figures S1A-S1B)**.

**Figure 1.**
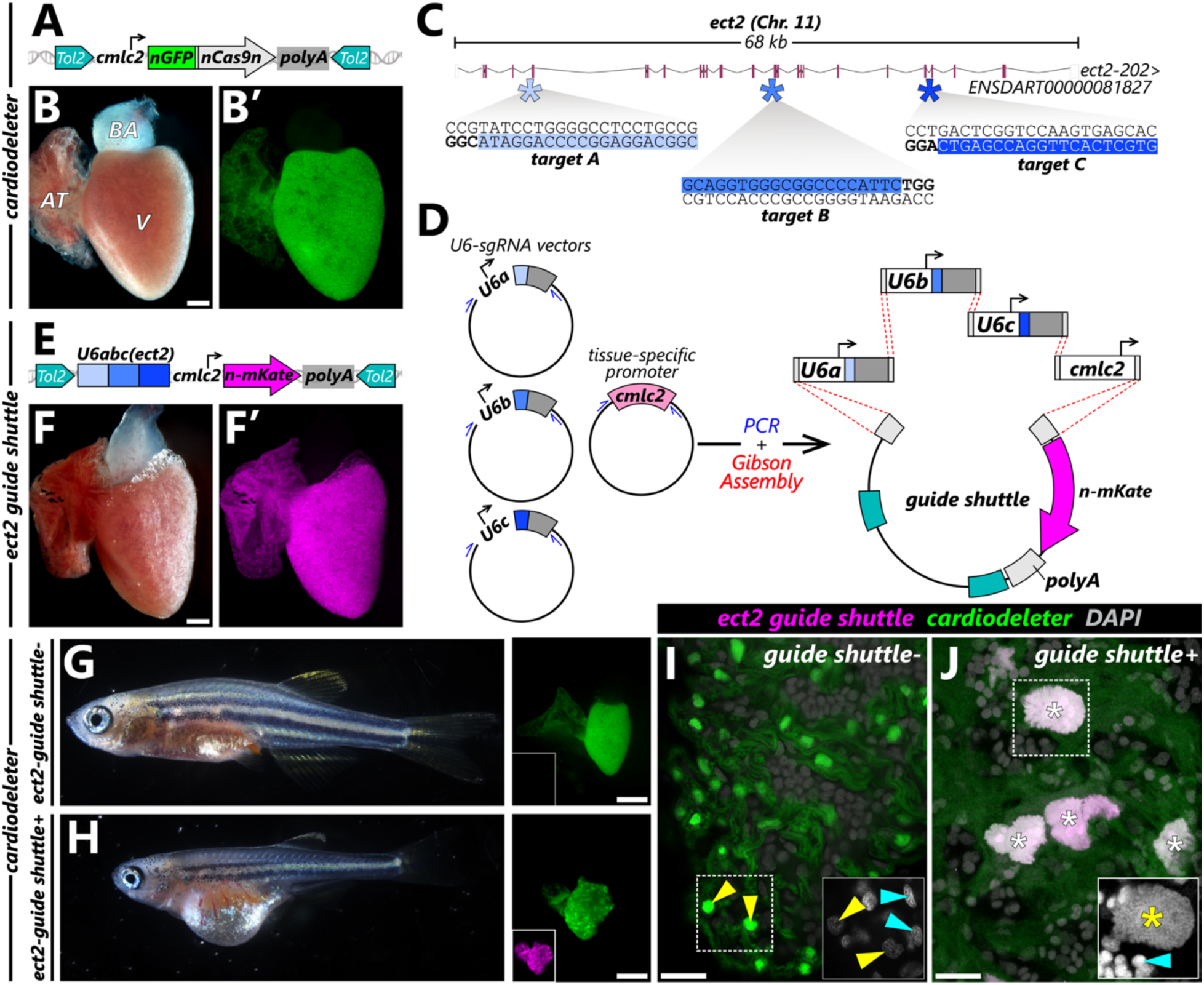
The *cardiodeleter* transgene efficiently creates cardiomyocyte-specific loss-of-function phenotypes in combination with gene-specific gRNAs. **(A)** Schematic representation of the *cardiodeleter* transgene, which drives the expression of nuclear GFP (nGFP) and a zebrafish codon optimized version of Cas9 in cardiomyocytes. **(B,B’)** Representative heart from an adult zebrafish carrying the *cardiodeleter* allele, exhibiting normal morphology (B) and expression of nGFP (B’). **(C)** Representation of the *ect2* locus in the zebrafish genome. Filled boxes represent coding exons, empty boxes indicate the 5’ and 3’ untranslated regions (UTRs). Asterisks indicate the location three selected gRNA targets, predicted using CRISPRScan. Blue areas indicate the target sequence. Sequences in bold highlight the PAM. **(D)** Strategy to generate a *guide shuttle* carrying three U6 promoters (U6a, U6b, U6c), each driving the expression of a gRNA targeting a gene of interest, and a transgenesis reporter. **(E)** Representation of the stable *ect2 guide shuttle* transgene. **(F, F’)** Representative heart from an adult zebrafish carrying the *ect2 guide shuttle* exhibiting normal morphology (F) and expression of n-mKate (F’). **(G,H)** 30-dpf *cardiodeleter*^+^ (G) and *cardiodeleter^+^ect2 guide shuttle^+^* (H) animals. Stalled growth and edema can be observed only in double transgenic animals. Dissected hearts are shown in the right panels. **(I,J)** Confocal images of DAPI-stained whole hearts from 30-dpf *cardiodeleter*^+^ (K) and *cardiodeleter^+^ect2 guide shuttle^+^* (L) animals. Boxed regions show DAPI staining at higher magnification (white). Asterisks: polyploid nuclei. Cyan and yellow arrowheads: non-myocytes and cardiomyocytes, respectively. AT, atrium; BA, bulbus arteriosus; Chr, chromosome; V, ventricle. Scale bars: 200 μm (B, F) and 20 μm (I, J).

Before using this system as the springboard for optimizing cardiomyocyte-specific *crispant* conditions, we first focused on determining whether our *cardiodeleter* strain efficiently induces mutations in targeted genes. To this end, we established an independent transgene driving the expression of three gRNAs targeting the cytokinesis regulator *ect2*. We chose this target because we have previously demonstrated that cells lacking *ect2* fail cytokinesis and become polyploid^27^, a phenotype we can detect based on nuclear size. Importantly, this phenotype emerges exclusively in homozygous mutant cells, meaning only biallelic mutations would result in cardiomyocyte polyploidization. Using CRISPRScan^28^, we selected three gRNAs without predicted off-targets in the zebrafish genome **(Figure 1C)**. We cloned these gRNA targets in vectors containing the *U6a*, *U6b*, and *U6c* zebrafish promoters^22^. Then, we devised a PCR and Gibson Assembly-based strategy to generate a *Tol2* transposable element containing these three U6-gRNA cassettes (**Figure 1D**, see Extended Protocols). This construct, which we call *guide shuttle*, contains the instructions for making the gRNAs and a transgenesis reporter, which labels cells of interest that integrated the vector. For our purposes, we chose the *cmlc2* promoter to drive the expression of the red fluorescent protein mKate in cardiomyocyte nuclei. As expected, animals carrying the *ect2 guide shuttle* (hereafter, *ect2-gs^+^*) developed normally to adulthood, exhibited strong mKate fluorescence in cardiomyocytes **(Figures 1E-1F)**, and showed normal cardiac histology **(Figure S1A)**. Thus, the independent expression of either Cas9 or *ect2* gRNAs does not induce aberrant phenotypes that could preclude using these lines in future studies.

To determine whether the co-expression of Cas9 and gRNAs targeting *ect2* induces biallelic mutations that result in cardiomyocyte polyploidization, we crossed adult animals carrying each of these transgenes. Double transgenic animals were externally indistinguishable from single transgenics at larval stages. However, at around 30 days post-fertilization (dpf), *cardiodeleter^+^ect2-gs^+^*animals exhibited stalled growth, edema, and abnormal cardiac morphology **(Figures 1G-1H)**. Analysis of their hearts revealed myocardial polyploidization, evidenced by a reduced number of cardiomyocyte nuclei and increased nuclear size **(Figures 1I-1J)**. This phenotype is consistent with our previous results inducing cytokinesis failure in cardiomyocytes using a dominant negative version of *ect2* (*cmlc2:dnEct2*, ref.^27^). As described for the *cmlc2:dnEct2* animals, *cardiodeleter^+^ect2-gs^+^* animals developed a highly penetrant heart failure phenotype by 45 dpf. Consequently, we recovered less than ∼4% of double-transgenic adults (**Figure S1A,** 3/90 animals), while most *cardiodeleter^+^ect2-gs^−^* siblings (86/90) survived to adulthood.

The myocardial polyploidization phenotype suggests that our strategy efficiently disrupts the *ect2* locus. To identify mutations generated by Cas9 in this gene, we extracted genomic DNA from double-transgenic animals at 30 dpf, when the polyploid phenotype was evident **(Figure 1H)**. We sequenced a small amplicon that encompasses guide A-target, predicted to be our most efficient gRNA (CRISPRScan score=85, *vs.* 62 and 65 from guides B and C, respectively). Sequencing revealed the presence of mutant variants in the sample **(Figures S1C-S1D)**, albeit occurring at a very low frequency (∼1.4% of 73,579 Next-Generation Sequencing reads). Given that our *ect2-gs* drives the expression of 3 gRNAs targeting distant genomic sequences, Cas9 may generate large deletions within the *ect2* locus. To test this hypothesis, we mapped three potential deletion alleles arising from the excision of the intervening sequences flanked by each pair of gRNA targets. We designated these deletion (Δ) alleles as ΔAB, ΔAC, and ΔBC and devised a PCR-based strategy for their detection **(Figure S1C)**. We successfully amplified each of the predicted Δ alleles in DNA extracted from *cardiodeleter^+^ect2-gs^+^* hearts but failed to obtain amplification from the fins of the same animals **(Figure S1E)**, confirming the presence of large deletions specifically in the heart. Sequencing of the Δ amplicons further confirmed the presence of the predicted deletions **(Figure S1F)**. These results demonstrate that our *cardiodeleter* system efficiently disrupts targeted genes in cardiomyocytes when combined with a *guide shuttle* that drives the expression of the corresponding gRNAs. Thus, the *cardiodeleter* is a versatile platform for refining various parameters to generate cardiomyocyte-specific *crispants*.

### An orthogonal transposase approach is required for the mobilization of the guide shuttle without disrupting the cardiodeleter insertion

The strategy described above to generate zebrafish that lack *ect2* in cardiomyocytes requires breeding stable lines and is, therefore, slow and time-consuming. To accelerate the generation of cardiomyocyte-specific *crispants*, we compared the efficiency of two approaches to generate biallelic mutations: injecting either synthetic gRNAs or a *guide shuttle* encoding the same gRNAs into *cardiodeleter^+^*embryos at the one-cell stage. Although these strategies differ in throughput and labeling of susceptible cells, both approaches could bypass the need to generate stable insertion for analysis, significantly accelerating this pipeline.

Injecting synthetic gRNAs into embryos carrying a tissue-specific Cas9 line has been reported as an efficient method to induce tissue-specific mutations. Specifically, this method has been successfully exploited in combination with neutrophil and macrophage-specific Cas9 lines^29,30^. Although the mutant cells remain unlabeled, the main advantage of this method is that it does not involve any cloning, increasing throughput. However, given that the injected gRNAs degrade over time, the efficiency of this approach may be suboptimal if the tissue-specific promoter driving Cas9 activates later in development. To test whether the delivery of synthetic gRNAs into *cardiodeleter+* embryos efficiently induces myocardial mutagenesis, we chose to target *tnnt2a*, which encodes for a cardiac troponin that can be recognized using a well-characterized antibody^31^ **(Figure 2A)**. We reasoned that biallelic mutations in *tnnt2a* would result in loss of antibody staining, allowing us to easily assess and quantify the efficiency of our system on a per-cell basis. To test whether the selected gRNAs were efficient at inducing biallelic mutations, we first co-injected the gRNAs and Cas9 protein at the one-cell stage. As predicted, this approach generated global *crispants* with a high percentage of cardiomyocytes that lacked Tnnt2a expression, validating our gRNAs and experimental approach **(Figures 2B-2C)**. While providing Cas9 at the one-cell stage was highly efficient, injecting the same gRNAs into *cardiodeleter+* embryos resulted in no detectable loss of Tnnt2a staining **(Figures 2D-2E)**. These results suggest that by the developmental stage that the *cardiodeleter* transgene activates, the gRNAs are either entirely degraded or present at insufficient levels to induce biallelic mutations.

**Figure 2.**
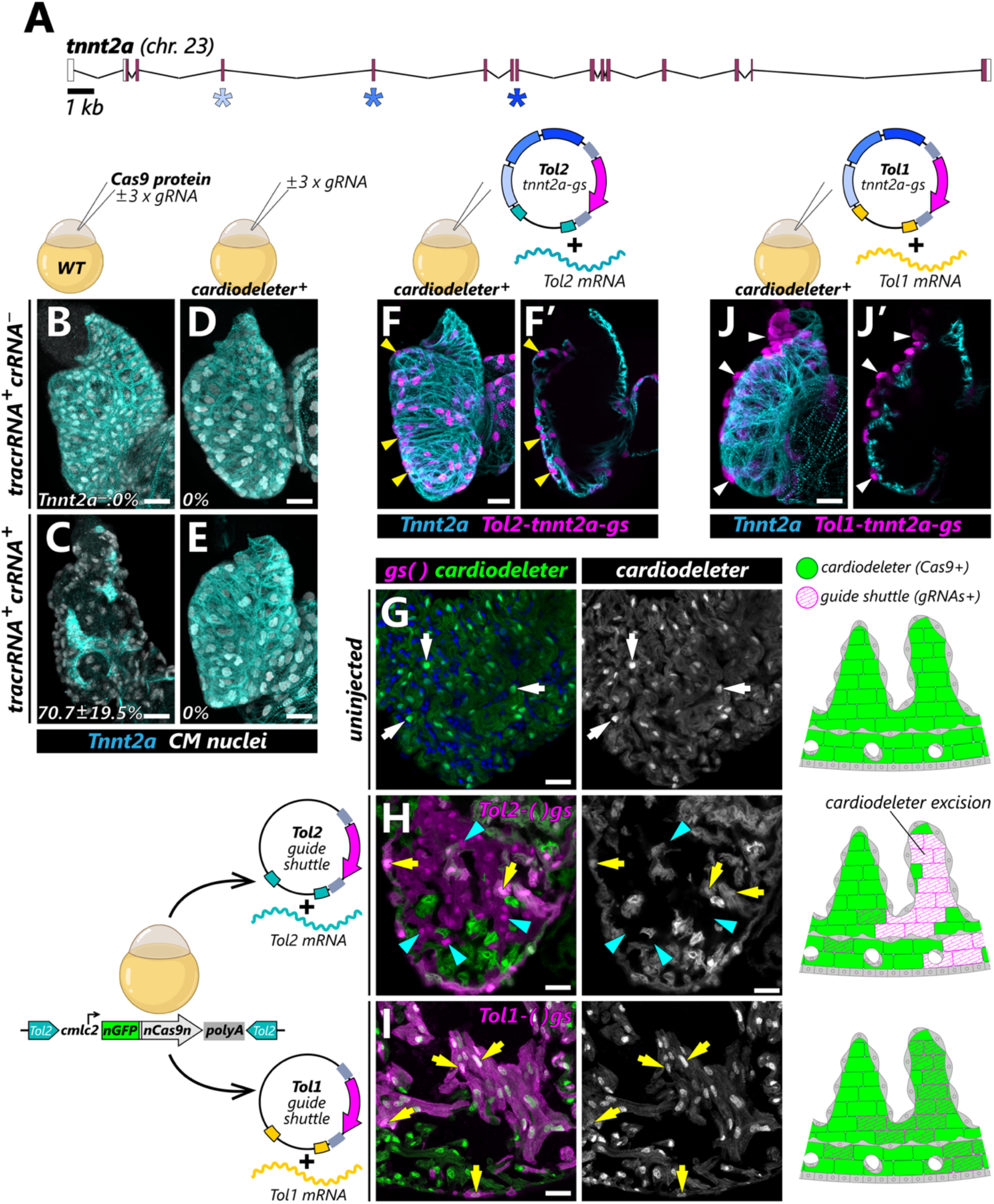
An orthogonal transposase approach is required to efficiently generate cardiomyocyte-specific *crispants*. **(A)** Representation of the *tnnt2a* locus in the zebrafish genome, indicating the exons targeted by the selected gRNAs (asterisks). **(B-E)** Confocal projections of representative 96 hpf embryonic hearts from wild-type (B-C) or *cardiodeleter^+^*animals (D-E) from the indicated experimental groups, immunostained to detect cardiomyocyte nuclei (white) and Tnnt2a (cyan). The percentage of Tnnt2a-negative (Tnnt2a^−^) cardiomyocytes is summarized in the bottom left corner (average ± standard deviation, *n* = 4, 7, 6, and 8 embryos). **(F,J)** Confocal projections of 96 hpf embryonic hearts from animals carrying the *cardiodeleter* transgene and injected with a *Tol2-* (F, n=8) or *Tol1*-(J, n=7) based guide shuttle encoding guide RNAs targeting *tnnt2a*, immunostained to detect Tnnt2a (cyan). Yellow and white arrowheads indicate Tnnt2a^+^ and Tnnt2a^−^ *guide shuttle^+^*cardiomyocytes (magenta), respectively. Single confocal planes are shown in F’ and J’. **(G-I)** Representative ventricular sections from ∼30 dpf *cardiodeleter^+^* animals, uninjected (G) or injected with the indicated combinations of *guide shuttles* and mRNA (H, I). White arrows indicate cardiomyocyte nuclei. Yellow arrows and cyan arrowheads indicate mKate^+^*cardiodeleter^+^*and mKate^+^*cardiodeleter^−^* cardiomyocytes, respectively. Cartoons (right) summarize the observed patterns of excision and integration of the cardiodeleter and guide shuttles. Chr, chromosome. Scale bars: 20 μm.

To achieve a more sustained expression of the gRNAs, we assembled a *guide shuttle* encoding the same guides targeting *tnnt2a* (*tnnt2a-gs)* and co-injected it into *cardiodeleter*+ embryos at the one-cell stage with Tol2 mRNA. While assembling a *guide shuttle* requires cloning and reduces throughput, this method ensures a continuous expression of the guides and enables rapid identification of the presumptively mutant cells. Despite the conceptual advantages of this approach, previous attempts to mobilize a construct containing a *U6* promoter driving the expression of a gRNA into tissue-specific Cas9 lines showed extremely low mutagenesis efficiency, ∼43 times lower than the levels achieved via synthetic gRNA injection^29^. Consistent with these results, we found no evidence of biallelic editing in *cardiodeleter*+ embryos injected with the *tnnt2a-gs*, despite efficiently incorporating the *guide shuttle* into cardiomyocytes **(Figures 2F-2F’)**.

A necessary condition for successfully generating tissue-specific *crispants* is efficiently mobilizing the *guide shuttle* to the genome without affecting the *cardiodeleter* transgene. Because both our Cas9 transgene and the *guide shuttle* are Tol2-based constructs, we examined whether the injection of *Tol2* mRNA for transposition of the *guide shuttle* was excising the *cardiodeleter*. To test this, we injected an empty version of our Tol2-based *guide shuttle* into *cardiodeleter* embryos and analyzed the expression of the nGFP-Cas9 cassette. Uninjected animals showed uniform expression of nGFP-Cas9 across the myocardium **(Figure 2G)**. In contrast, we detected large nGFP-Cas9 negative clones in animals injected with Tol2 mRNA and the Tol2-based *guide shuttle* **(Figure 2H)**, indicating excision of the *cardiodeleter* transgene. Thus, injecting *Tol2*-based *guide shuttles* directly into *cardiodeleter* animals is not a viable strategy to generate cardiomyocyte-specific *crispants*.

To circumvent this limitation, we sought to replace the *Tol2* sites in our *guide shuttles* with sequences that will be specifically recognized by a different transposase. We chose the *Tol1* transposase from medaka, which efficiently mobilizes *Tol1*-based elements but fails to recognize *Tol2*-specific sequences^32^. An additional advantage of this system is that there is an available zebrafish codon-optimized version of this transposase^33^. Co-injection of a *Tol1*-based *guide shuttle* and *Tol1* mRNA into *cardiodeleter* embryos resulted in efficient transposition of the empty *guide shuttle* without losing the *cardiodeleter* transgene **(Figure 2I)**. These results suggest that this orthogonal transposase approach could be useful in delivering the guides to cells expressing Cas9 from a Tol2-based transgenic insertion. To further test our approach, we generated a *Tol1*-based *tnnt2-gs* and injected it into *cardiodeleter* embryos at the one-cell stage. In contrast to the *Tol2-tnnt2-gs*, we detected Tnnt2a^−^ cardiomyocytes in *cardiodeleter* animals injected with the *Tol1-tnnt2-gs* **(Figures 2J-2J’)**. Consistent with models of transplantation of cells lacking other sarcomeric proteins^34,35^, Tnnt2a^−^ cardiomyocytes bulged outwards as individual cell aneurysms without affecting wild-type cardiomyocytes. Collectively, these results demonstrate that *Tol1*-based *guide shuttles* can be efficiently mobilized into the genome without affecting Tol2-based transgenes.

### Labeled cardiomyocyte clones carrying biallelic mutations in sarcomeric genes survive to adulthood intermingled with wild-type cells

We next aimed to test to what extent our cardiomyocyte-specific *crispant* approach successfully induces LOF phenotypes that (1) are efficient and restricted to the population labeled by the *guide shuttle* reporter, (2) can be detected shortly after injection, (3) bypass the embryonic lethality that germline mutations would produce, and (4) affect genes located in different regions of the genome. To expand our analysis beyond the characterization of *tnnt2a*, we chose two additional targets that also encode structural proteins involved in cardiac contraction: *cmlc2* (official symbol, *myl7*), and *amhc* (official symbol, *myh6*) **(Figure 3A)**. Mutations in these genes, identified in ENU screens for cardiovascular phenotypes^2^, cause contractility defects and have been positionally cloned^36^. As for *tnnt2a*, there are available antibodies that specifically recognize the products of each of these genes^36,37^, facilitating the detection of biallelic mutations on a per-cardiomyocyte basis.

**Figure 3.**
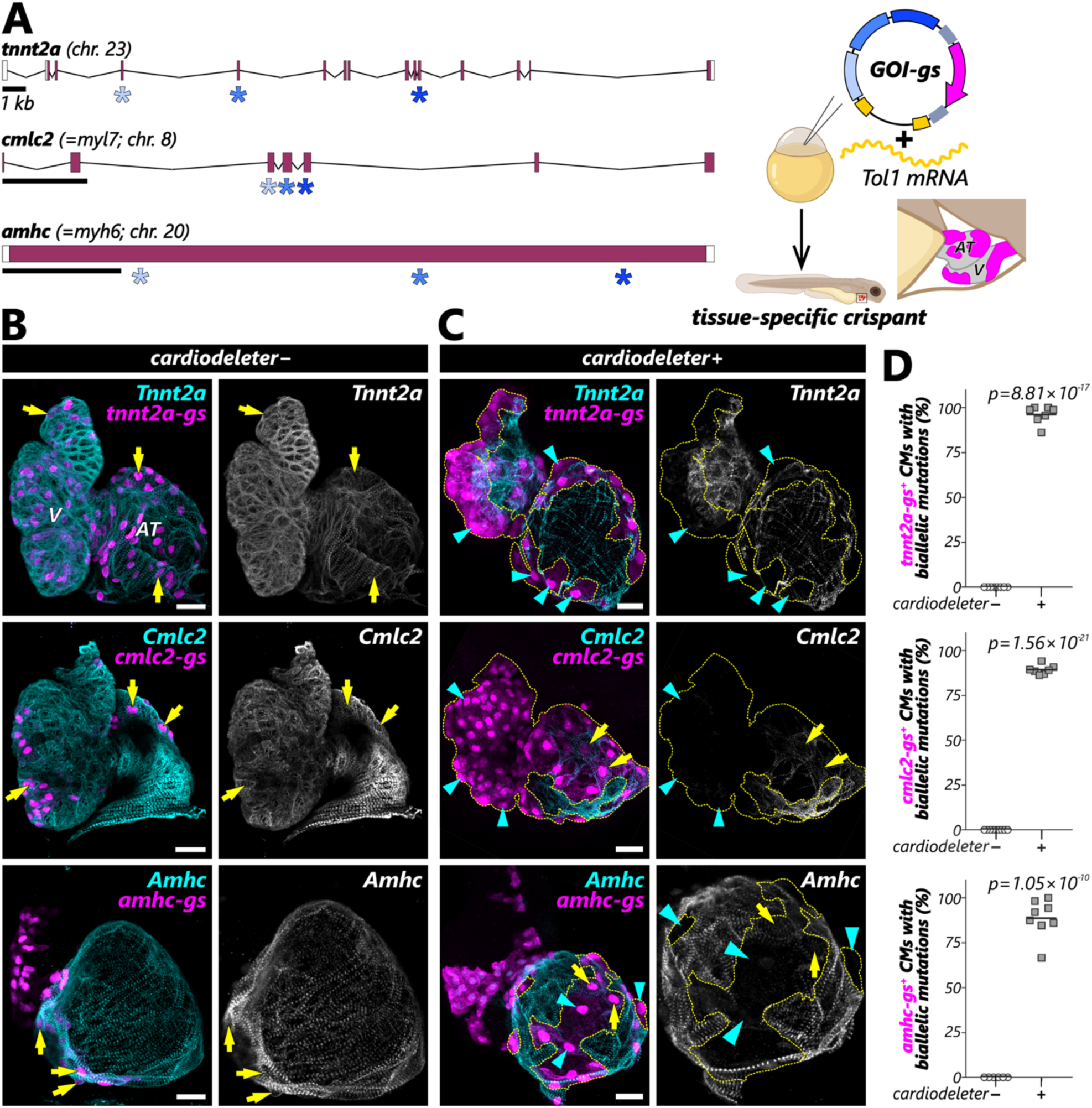
Cardiodeleter-induced *crispants* carry biallelic mutations in ∼90% of labeled cardiomyocytes. **(A)** Representation of the location of the gRNAs targeting *tnnt2a*, *cmlc2*, and *amhc*. Tol1-*guide shuttles* encoding three guides for each gene were injected into wild-type or *cardiodeleter^+^* embryos at the one-cell stage. Embryos exhibiting mKate^+^ clones in the heart, indicative of *guide shuttle* integration, were selected at 4 dpf and fixed for immunostaining. **(B,C)** Confocal projections of representative 96 hpf embryonic hearts from wild-type (B) or *cardiodeleter^+^* animals (C) injected with guide shuttles targeting the indicated genes, immunostained with specific antibodies to recognize the corresponding protein (cyan). Cyan arrowheads indicate *guide shuttle* (mKate)^+^ cardiomyocytes with biallelic mutations, leading to the complete loss of the corresponding protein. Yellow arrows indicate mKate^+^ cardiomyocytes maintaining protein expression. **(D)** Quantification of the efficiency of biallelic mutations in wild-type or *cardiodeleter* embryos injected with the indicated *guide shuttles*. Individual values plotted; black line: average. *p*-values: two-tailed unpaired t-test. AT, atrium; V, ventricle; chr, chromosome. Scale bars: 20 μm.

To target our selected genes, we first generated Tol1-based *guide shuttles* encoding three gRNAs and the transgenesis reporter. Next, we injected our *guide shuttles* into wild-type and *cardiodeleter+* embryos, fixed animals carrying mKate+ (gs+) clones at 4 dpf, and determined the presence of the target proteins using immunofluorescence **(Figure 3B)**. As expected, the expression of the gRNAs alone in wild-type animals did not affect the distribution of these proteins. By contrast, we detected patches of cardiomyocytes lacking the targeted proteins in *cardiodeleter*+ animals injected with the corresponding *guide shuttles* **(Figure 3C)**. Notably, the loss of the targeted proteins was restricted to mKate+ cells, indicating that the mutagenesis only happened in cells that incorporated the *guide shuttle*. To assess the efficiency of our method in inducing biallelic mutations, we next quantified the percentage of cells that incorporated the *guide shuttles,* and that lacked the expression of the corresponding protein. At 4 dpf, ∼90% of all mKate+ cardiomyocytes displayed no staining for the targeted protein (*tnnt2a*: 96.2% ± 4.9, *cmlc2*: 89.4% ± 2.6, *amhc*: 89.7% ± 9.6; mean ± SD; n=7 embryos/gene), suggesting that our system results in a highly efficient biallelic gene disruption **(Figure 3D)**. Thus, our tissue-specific *crispant* strategy efficiently generates labeled clones with biallelic mutations that can be detected just a few days after injection.

Homozygous mutations in *tnnt2a* and *cmlc2* result in embryonic lethality^35,38^, and only ∼30% of *amhc^-/-^* survive to adulthood^37^. By contrast, we did not observe increased mortality in our cardiac *crispant* larvae, suggesting that the mosaicism in these hearts is sufficient to bypass early mortality. To determine whether the labeled mutant cells persist in the heart intermingled with wild-type cardiomyocytes, we grew *cardiodeleter+* animals injected with each of our *guide shuttles* after selecting candidates with mKate+ clones at 72 hpf **(Fig. 4A)**. As expected, the labeled cells expanded during cardiac growth, and by 40 dpf had formed clusters of mKate+ cardiomyocytes covering large patches of the myocardium **(Figures 4B-4D’)**. Immunostaining in sections revealed that virtually all mKate+ cardiomyocytes were negative for the proteins encoded by the targeted genes **(Figures 4E-4G)**. In *tnnt2a* and *cmlc2* myocardial *crispants* labeled clones exhibited abnormal histological organization **(Figures 4E’-4G’)**. These adult phenotypes had never been documented before due to the embryonic lethality of these mutants. Notably, the myocardial defects in *amhc* myocardial *crispants* were limited to the atrium, consistent with the atrial-specific expression of this myosin and ruling out an unspecific phenotype induced by the action of Cas9 **(Figures 4G-4G’)**. Moreover, atrial Amhc^−^ clones lacked myosin heavy chain (MHC) proteins but no other sarcomeric elements, whereas ventricular clones maintained MHC expression **(Figure S2A)**, phenocopying the *amhc* (*weak atrium*) mutant^36^. Collectively, these results demonstrate that our strategy generates clones of mutant cardiomyocytes that can be recovered in the adult heart, intermingled with wild-type cells.

**Figure 4.**
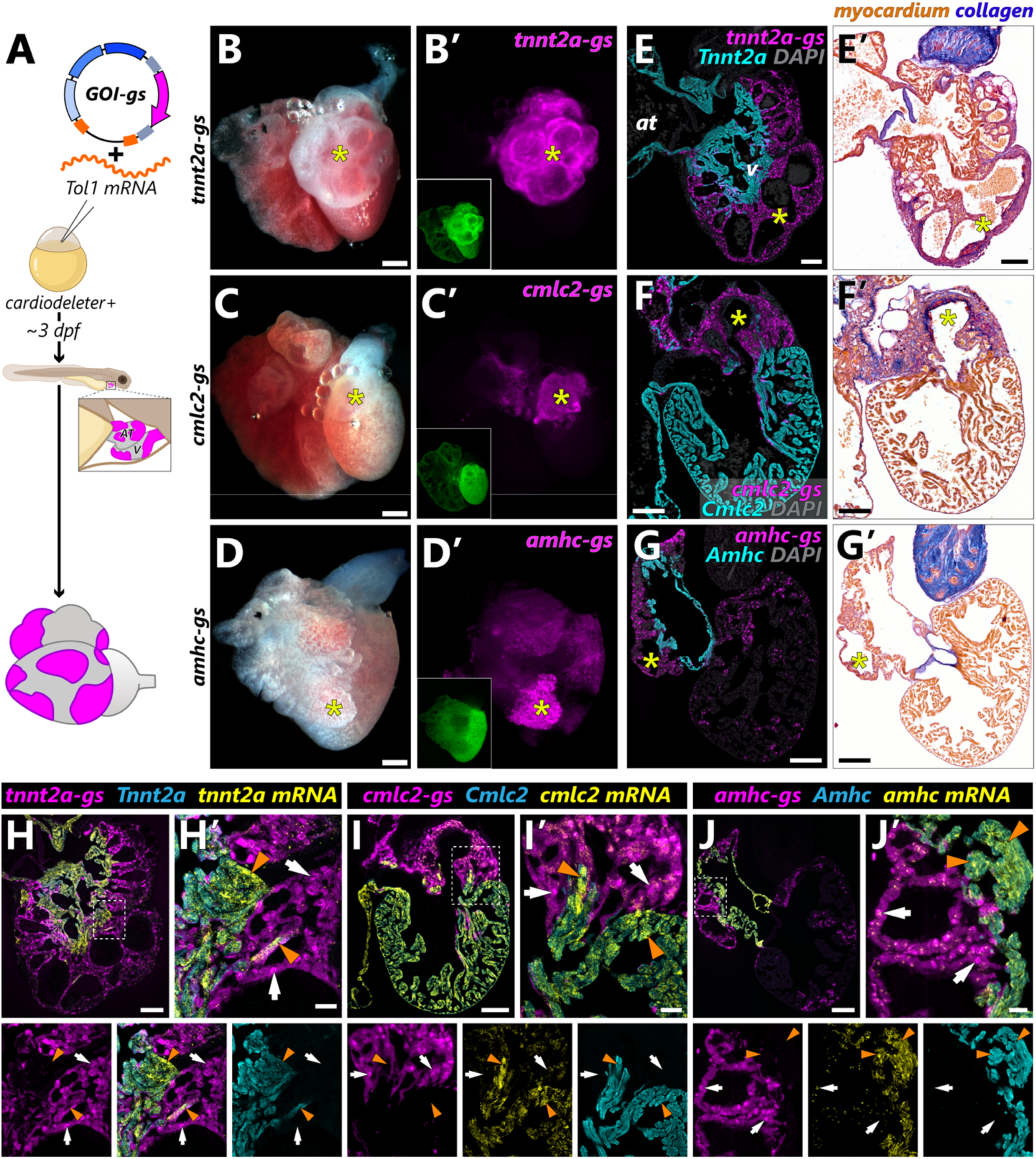
Cardiomyocyte clones carrying biallelic mutations in *tnnt2a*, *cmlc2*, or *amhc* survive to adulthood intermingled with wild-type cells. (A) Schematic representation of the experimental approach to test whether labeled myocardial clones carrying biallelic mutations can be recovered in adult hearts. **(B-D)** Representative images of dissected hearts from *cardiodeleter^+^*animals injected with the indicated *guide shuttles*. **(B’-D’)** mKate signal from hearts shown in B-D, indicating cardiomyocyte clones that have incorporated the corresponding *guide shuttle*. The GFP fluorescence from the *cardiodeleter* transgene is shown in the images located in the bottom left corner. **(E-G)** Histological sections from hearts shown in B-D, immunostained with antibodies to identify *guide shuttle^+^* cardiomyocytes (mKate^+^, magenta) and the corresponding protein encoded by the targeted gene (cyan). **(E’-G’)** Consecutive sections from hearts shown in E-G stained using the AFOG technique to identify collagen (blue) and myocardium (orange/brown). Yellow asterisks indicate large mKate^+^ clones that show complete absence of the corresponding protein. **(H-J)** Combined immunofluorescence-RNAScope in consecutive sections from hearts shown in E-G. Signal from mRNA ISH appears in yellow. White arrows indicate *guide shuttle* (mKate)^+^ cardiomyocytes with biallelic mutations, leading to the reduction of the corresponding mRNA and the complete loss of the corresponding protein. Orange arrowheads point to *guide shuttle*^−^ cardiomyocytes that maintain high levels of mRNA and protein expression. at: atrium; v, ventricle. Scale bars: 200 μm (B, C, D), 100 μm (E-I and E’-G’), and 20 μm (H’-J’).

Next, we tested whether we could detect changes in the mRNA levels of the targeted genes in our myocardial *crispants*. This strategy may be relevant for validation purposes when an antibody against the protein encoded by the GOI is unavailable. To detect potential changes in mRNA expression, we used RNAScope *in situ* hybridization in adjacent sections from 40 dpf myocardial *crispants*. We co-stained these sections with antibodies against the corresponding protein and mKate, which identifies cardiomyocytes carrying the *guide shuttle* **(Figures 4H-4J’)**. We detected a consistent reduction in mRNA levels for the three analyzed genes in cardiomyocytes carrying the *guide shuttle*, compared to the adjacent *gs^−^*myocardium (*tnnt2a*: 93.4% ± 3.5, *cmlc2*: 66.9% ± 6.6, *amhc*: 87.5% ± 2.8; relative reduction in expression levels; average ± SD; n=3 hearts/gene). These results demonstrate that, in the absence of specific antibodies, *in situ* hybridization can be used to validate efficient disruption of the genes targeted by the *cardiodeleter*.

### Highly efficient biallelic mutagenesis requires *guide shuttles* encoding multiple gRNAs

In our initial design of the *guide shuttles*, we included three gRNAs targeting our GOI. Although we reasoned that this approach would maximize mutagenesis, we sought to experimentally test whether reducing the number of gRNAs would significantly affect our ability to generate biallelic deletions. To this end, we assembled *guide shuttles* encoding three, two, or a single gRNA targeting *amhc* (*U6abc-*, *U6ab-*, and *U6a-amhc-gs*, respectively) and injected them into *cardiodeleter+* embryos at the one-cell stage. We focused on *amhc* because, in contrast to *ect2*, *tnnt2*, or *cmlc2*, ∼30% of the global *amhc* mutants survive to adulthood. This allows us to analyze the phenotypes emerging from F1 lines where the gene is deleted in all cardiomyocytes. Consistent with our previous findings, injection of *U6abc-amhc-gs* induced robust loss of Amhc staining in over 90% of mKate+ atrial cardiomyocytes at 4 dpf **(Figures 5A, 5D)**. The efficiency of biallelic deletion dropped to ∼70% and ∼37% in animals injected with *U6ab- and U6a-amhc-gs*, respectively **(Figures 5B-5D)**. These results suggest that the likelihood of inducing biallelic mutations decreases with each gRNA that is omitted.

**Figure 5.**
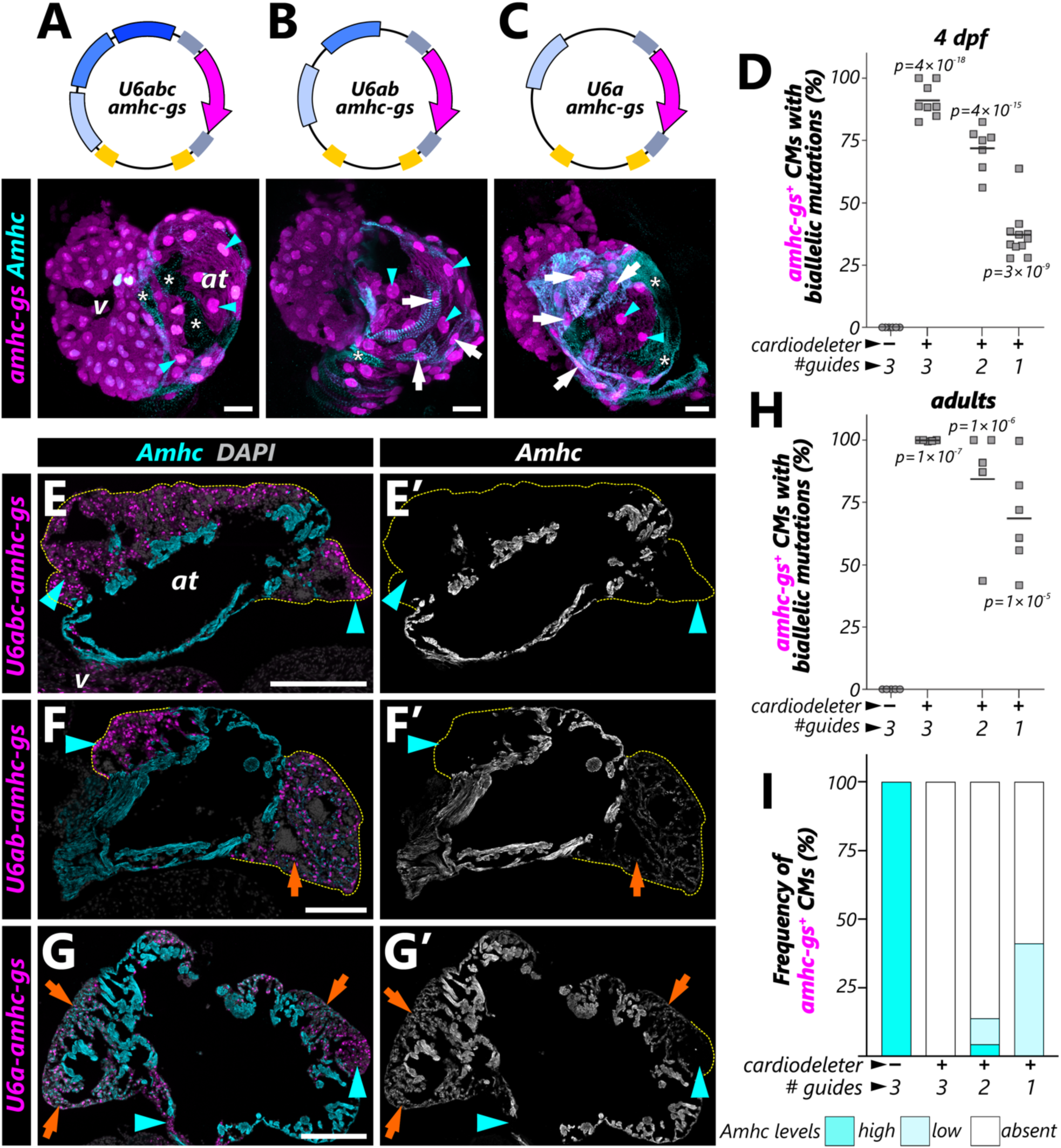
A *guide shuttle* encoding multiple gRNAs targeting *amhc* is required for highly efficient biallelic mutagenesis. **(A-C)** Confocal projections of representative 96 hpf embryonic hearts from *cardiodeleter^+^* animals injected with *guide shuttles* carrying three (A), two (B) or one (C) gRNAs targeting *amhc*. **(D)** Quantification of the efficiency of biallelic mutations in wild-type and *cardiodeleter* embryos injected with the indicated *guide shuttles*. Individual values plotted; black line: average. *p*-values: one-way ANOVA, followed by Tukey’s multiple comparisons test. **(E-G)** Histological sections of hearts from adult *cardiodeleter* animals injected with the indicated *guide shuttle*, immunostained with antibodies to identify mKate (*guide shuttle*, magenta) and Amhc (cyan). Amhc signal is shown in E’-G’. Cyan arrowheads indicate mKate (*guide shuttle*)^+^ cardiomyocytes with biallelic mutations that lead to complete depletion of the Amhc protein. Orange arrows indicate *guide shuttle*^+^ cardiomyocytes that maintain detectable levels of Amhc expression. **(H)** Efficiency of biallelic mutations in wild-type and *cardiodeleter* adult animals injected with the indicated *guide shuttles*. Individual values plotted; black line: average. *p*-values: one-way ANOVA, followed by Tukey’s multiple comparisons test. **(I)** Distribution of *guide shuttle*^+^ cardiomyocyte population according to Amhc protein levels in wild-type and *cardiodeleter* adult animals injected with the indicated *guide shuttles*. at: atrium; v, ventricle. Scale bars: 20 μm (A, B, C), 100 μm (E-G).

Next, we tested whether the efficiency of biallelic mutagenesis increases as a function of time, which could minimize the observed differences derived from using less than three guides. To this end, we injected each *amhc*-*guide shuttle* described above and analyzed *crispants* at 40 dpf. We found that the efficiency of biallelic mutagenesis was consistently higher at this stage than in the embryo. Virtually all mKate+ cardiomyocytes were Amhc*^−^*when the *guide shuttle* encoded 3 gRNAs (99.8%, **Figures 5E, 5H**). In contrast, the efficiency of biallelic mutagenesis increased to ∼84% and ∼68% when using two guides and a single gRNA, respectively **(Figures 5F-5H)**. Importantly, while the results using three gRNAs were consistent across animals, we detected variability in the frequency of biallelic deletions in animals injected with *U6ab-* or *U6a-amhc-gs*, ranging from ∼40% to ∼100% **(Figure 5H)**. Decreasing the number of gRNAs resulted in *amhc-gs^+^* cells that retained Amhc staining, indicating monoallelic deletions that reduce but do not eliminate Amhc expression **(Figures 5F’, 5G’, and 5I)**. This phenotype was particularly noticeable when using a single gRNA, where over 40% of mKate+ cardiomyocytes retained detectable levels of Amhc **(Figure 5G, 5G’)**.

We next sought to determine whether there were differences in efficiency in biallelic deletion between *crispants* and stable lines carrying each of the versions of our *amhc-gs*. To this end, we identified founders injected with each construct, crossed them with *cardiodeleter+* animals, and raised double transgenics. Consistent with the original description of the *amhc* mutant, ∼70% of our *cardiodeleter^+^U6abc-amhc-gs^+^* embryos died during the first week of life due to atrial contractility defects. Surviving double transgenic adults phenocopied the severe atrial hypoplasia described in the mutant^39^, and all mKate+ atrial cardiomyocytes were Amhc*^−^* **(Figures S2B, S2C)**. In contrast, the efficiency of biallelic mutagenesis dropped to ∼85% and ∼56% when the *guide shuttle* encoded two guides or a single gRNA, respectively **(Figures S2B, S2C)**. Thus, the loss of Amhc immunoreactivity and the degree of atrial hypoplasia appeared proportional to the number of gRNAs encoded by the *guide shuttle*. Our results indicate that the efficiency of biallelic deletions in adult *crispants* is an excellent proxy for that of stable transgenics carrying the *guide shuttle*. Furthermore, we concluded that, when technically feasible, using a *guide shuttle* encoding three gRNAs in combination with the *cardiodeleter* maximizes the likelihood of generating biallelic mutations.

### Long-term tracking of *erbb2* myocardial *crispants* and stable-myocardial mutants demonstrates a cell-autonomous role in trabeculation

Recovering adult hearts with mutant clones prompted us to consider whether our approach could replace blastomere transplants to perform cell autonomous studies. Although very informative, blastomere transplantations must be analyzed before the host rejects the grafted cells, hindering the analysis of the long-term contributions of the mutant cells to the adult organ. Thus, we next sought to test whether we could exploit our approach as a replacement for the classical cell transplantation assays to analyze adult phenotypes. As a proof of concept, we focused on *errb2*, a gene essential for ventricular trabeculation in zebrafish^23^. When mutant blastomeres are transplanted into wild-type embryos, *erbb2*^-/-^ cardiomyocytes contribute exclusively to the primordial myocardium^23^. However, due to the limitations of the transplantation approach, the contributions of these mutant cardiomyocytes beyond 5 dpf remain unknown. To disrupt *errb2* in cardiomyocytes, we assembled a *guide-shuttle* encoding three guides targeting *erbb2* **(Figure 6A)** and injected it into WT and cardiodeleter+ embryos at the one-cell stage. Consistent with well-established phenotypes, cardiomyocytes lacking *erbb2* (*cardiodeleter+ erbb2-gs+*) contributed exclusively to the primordial layer at 5 dpf. Trabecular cardiomyocytes were derived exclusively from *erbb2-gs*^−^ cells **(Figures 6B, 6C)**. While *erbb2-gs+* cardiomyocytes contributed to both the primordial and trabecular myocardium in WT hearts at 70 dpf, we found restricted contribution to the primordial myocardium in *cardiodeleter+* animals **(Figure 6D-6E’’)**. These results support the notion that *erbb2* is cell-autonomously required in cardiomyocytes to give rise to the trabecules and demonstrate that our experimental approach circumvents previous limitations of the blastomere transplant paradigm.

**Figure 6.**
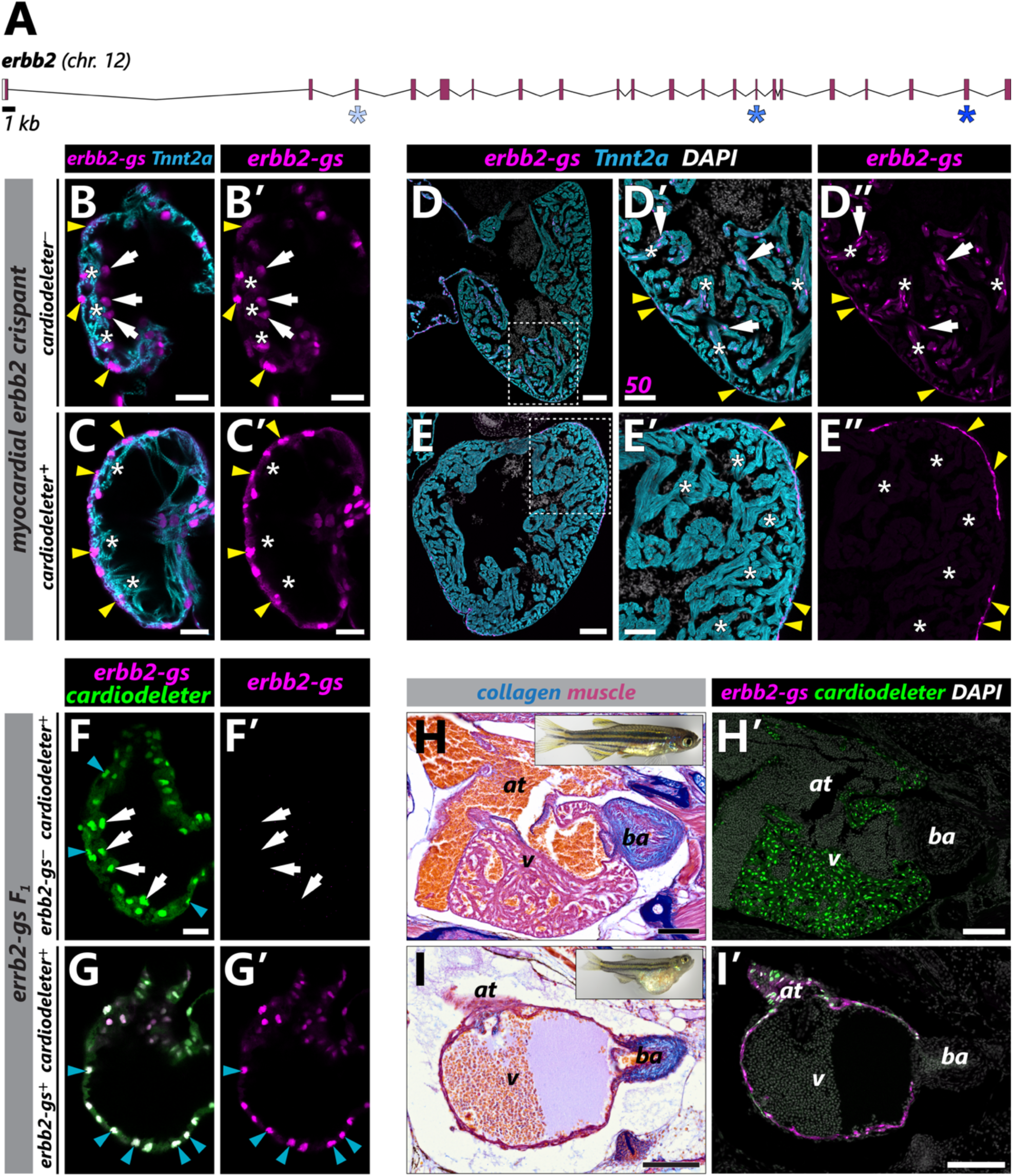
Long-term tracking of *erbb2* myocardial *crispants* and stable-myocardial mutants demonstrates a cell-autonomous role in trabeculation. (A) Representation of the location of the gRNAs targeting *erbb2*. **(B,C)** Representative single plane confocal images of embryonic hearts at 4 dpf from wild-type (B) and cardiodeleter+ animals (C), injected with a *guide shuttle* encoding gRNAs targeting *erbb2 (erbb2-gs)*, immunostained to detect Tnnt2a (cyan). **(D,E)** Representative heart sections from animals of the indicated genotypes, injected with the erbb2-gs, analyzed at 40 dpf. Magnifications of the boxed areas in D and E are shown in D’-E’’. Yellow arrowheads: *guide shuttle*^+^ cardiomyocytes located at the primordial myocardium; white arrows: *guide shuttle*^+^ trabecular cardiomyocytes; asterisks: trabecules. **(F, G)** Representative images of 4 dpf embryonic hearts from the indicated genotypes, immunostained to detect nGFP (*cardiodeleter*, green) and mKate (*erbb2-gs*, magenta). Cyan arrowheads: primordial cardiomyocytes; white arrows: trabecular myocardium. **(H,I)** Representative sections of *cardiodeleter*^+^ (H) and *cardiodeleter*^+^*erbb2-gs*^+^ siblings (I) stained with the AFOG technique to detect collagen (blue) and myocardium (orange/brown). Whole fish pictures shown on the top, right corner. **(H’, I’)** Consecutive sections from the hearts shown in H and I, immunostained to detect nGFP (*cardiodeleter*, green) and mKate (*erbb2-gs*, magenta). at: atrium; v, ventricle. Scale bars: 200 μm (B, C, D), 100 μm (E-I and E’-G’), and 20 μm (H’-J’).

Zebrafish deficient for *erbb2* die at larval stages, exhibiting defects in the cardiovascular system and absence of myelination of the peripheral nervous system^40,41^. Taking advantage of the tissue-specificity of our system, we next aimed to determine the cardiac phenotype emerging from the loss of *erbb2* exclusively in the myocardium. To this end, we identified animals stably transmitting the *erbb2-gs* to their progeny and crossed them with *cardiodeleter+* zebrafish. In contrast to their *erbb2-gs*^−^ siblings, double transgenic embryos exhibited no trabecular development at 5 dpf **(Figures 6F-6G’)**. Double transgenics grew initially without external defects but showed signs of heart failure by ∼30 dpf, including pericardial edema and stalled growth. A histological analysis of these animals revealed the complete absence of trabecular myocardium in double transgenic animals. Importantly, 100% of double transgenics developed this phenotype, and we could not recover animals beyond this stage, highlighting our system’s high efficiency and the phenotype’s penetrance.

## DISCUSSION

The zebrafish community has traditionally lacked robust approaches for tissue-specific reverse genetics, which have been successfully exploited in other model organisms for decades. For example, the Gal4/RNAi system, a cornerstone of *Drosophila* genetics^42^, has never been broadly used in zebrafish. Precise knock-ins, which have been part of the mouse genetics toolbox for many decades, have only recently been implemented in zebrafish^13^. Recent advances have made the generation of lox-flanked exons possible and successfully demonstrated the Cre-mediated elimination of one gene in a specific cell type^14,43^. Although these methods are the gold standard for precisely dissecting the role of a gene in a given tissue, generating these lines requires significant expertise and is extremely time-consuming, severely reducing throughput.

Somatic Cas9-mediated mutagenesis has emerged as a powerful tool for reverse genetics amenable to moderate and large-scale genetic screens^44,45^. *In vivo*, these methods entail delivering gRNAs to animals that stably express Cas9 in a tissue of interest, usually using viruses or nanoparticles^46^. This modularity represents a key advantage of the approach, as the same Cas9 transgene can be used to target virtually any gene. For the study of mammalian cardiomyocyte biology, myocardial-Cas9 lines have been established and characterized, and these strains are now widely used in the field^17,19,45,47,48^. Although these strategies are now commonly used in mice, the equivalent modular approach has not been developed for zebrafish. Here, we describe a cardiomyocyte-specific Cas9 transgene, the *cardiodeleter*, that can be modularly combined with different gRNAs to induce targeted mutagenesis. While mobilizing transposons encoding gRNAs into zebrafish Cas9-stable lines had proven highly inefficient^29^, we developed an orthogonal transposase approach that circumvents this limitation. Our *Tol1*-based *guide shuttles* enable long-term tracking of the presumptively mutant cells, which, combined with the high efficiency of biallelic mutations, creates the perfect model to develop myocardial-specific *crispants*. Using this intersection-based approach, only cells that express both Cas9 and the *guide shuttle*, easily identifiable based on the expression of fluorescent proteins, will carry mutations, while the neighboring cells can serve as the control. Although we chose the Tol1 system for our *guide shuttles*, other transposases that do not excise the Tol2-based *cardiodeleter* construct would be potential alternatives.

Delivering synthetic gRNAs into *cardiodeleter*+ embryos at the one-cell stage did not result in biallelic mutagenesis. This result contrasts with previous descriptions of other tissue-specific Cas9 alleles that generate high rates of mutagenesis^29,30^, but is consistent with the notion that both Cas9 and the gRNAs must simultaneously be present to induce mutations. Because the injected gRNAs are progressively degraded and diluted with each cell division, it is unlikely that they will be present by the time the *cardiodeleter* is robustly expressed. Although injecting synthetic gRNAs into tissue-specific Cas9 transgenes that are activated early in development may be a viable approach, the main limitation of this strategy is that the cells susceptible to carrying mutations remain unlabeled. Our *guide shuttle* approach solves these limitations: the information to continuously produce the gRNAs is stably inserted in targeted cells, which can be identified based on the expression of a fluorescent protein. Although most of our *guide shuttles* are designed to illuminate cardiomyocytes specifically, we have also generated alternative versions using the *ubiquitin b* (*ubb*) promoter, ensuring ubiquitous labeling of all cells that incorporate the construct (see Extended Protocols).

Most approaches to calculating mutagenesis’ efficiency using somatic CRISPR use amplicon sequencing. However, this approach does not allow researchers to determine whether an individual cell carries biallelic mutations. We targeted genes that encode proteins we can detect using specific antibodies for validation purposes. The rate of biallelic mutations using the *cardiodeleter* in combination with *guide shuttles* encoding three gRNAs is very high, with virtually all *guide shuttle*+ cells in adults showing complete depletion of the corresponding protein. We anticipate that the levels of Cas9 expression may influence the system’s efficiency, with weak promoters resulting in inconsistent editing. Additionally, we found experimental evidence that the number of gRNAs targeting a gene is an important variable that determines biallelic mutagenesis efficiency. CRISPR mutagenesis often activates the non-homologous end joining (NHEJ) pathway, which results in widely heterogeneous mutations. Consequently, lesions induced by Cas9 in a single location may lead to small in-frame insertions or deletions. Using more than one gRNA resulted in large deletions of the intervening sequences, maximizing the chances of inactivating the targeted gene. A potential alternative to reduce the unpredictability of the mutagenesis using a single guide would be to focus on gRNAs that are predicted to result in microhomology-mediated end joining (MMEJ). These gRNAs lead to a predictable mutant allele with high frequency. A dedicated, publicly available algorithm, MENTHU, could be exploited to identify these gRNAs^49^. To facilitate the dissemination of these technologies in the field, we have compiled a comprehensive protocol for guide identification, cloning in U6-containing plasmids, and assembly into the final Tol1 constructs (see Expanded Protocols). By standardizing the protocols and providing examples, our goal is to enhance the reproducibility of these approaches in any laboratory.

Despite its many advantages, we are aware of the limitations of our system. As with any other tissue-specific Cas9 lines, the *cardiodeleter* drives continuous expression of this nuclease from the time of myocardial specification. Although we have not found any evidence of cardiac toxicity in our system, we lack temporal control of the mutagenesis activity of Cas9. This strategy limits the use of the *cardiodeleter* in adult regeneration studies, where the absence of a given gene may result in developmental defects. We are validating new inducible systems, including a doxycycline-responsive Cas9 allele and a line expressing Cas9 in cardiomyocytes upon Cre-mediated recombination. Additionally, it may be impossible to find unique Cas9 targets in a GOI. Cas12/Cpf1 has become an alternative nuclease well-validated in zebrafish *crispant* approaches^50^. Establishing tissue-specific Cpf1 lines and validating whether we can efficiently deliver the appropriate gRNAs using *guide shuttles* would circumvent these limitations. Importantly, the methods we have established for the efficient delivery of multiple gRNAs with permanent labeling could be applied beyond the classical Cas9-mediated mutagenesis, including promoting and repressing transcription using deactivated forms of Cas9, which are still not broadly employed in zebrafish.

## Supporting information

Extended Protocols - Generation of gene-specific guide shuttles.

## Acknowledgments

J.M.G.-R. thanks J. Butler, B. Howell and the organizers of the 2024 Boston College Intersections Villa Faculty Writing Retreat for the opportunity to complete significant portions of this manuscript; Jesús Torrez-Vazquez (NYU) for helpful discussions; the Boston College Animal Care Facility and past and present members of the González-Rosa Lab for fish care and assistance; and Bret Judson of the Boston College Imaging Facility for training and assistance in image acquisition. DB received support from a Wallonie-Bruselles International Postdoctoral Fellowship. NJ was supported by a Summer Research Trainee Program (SRTP) Fellowship from the Massachusetts General Hospital Center of Diversity and Inclusion. LP received funding from the Vice Provost of Research Office of Boston College. Work in the Gonzalez-Rosa Lab is supported by the National Institutes of Health (R01HL164749), the American Heart Association (19CDA34660207), the Corrigan Minehan Foundation (SPARK Award), the Hassenfeld Foundation (Hassenfeld Research Scholar), and internal funds from Boston College.

## MATERIALS AND METHODS

### Key resources table

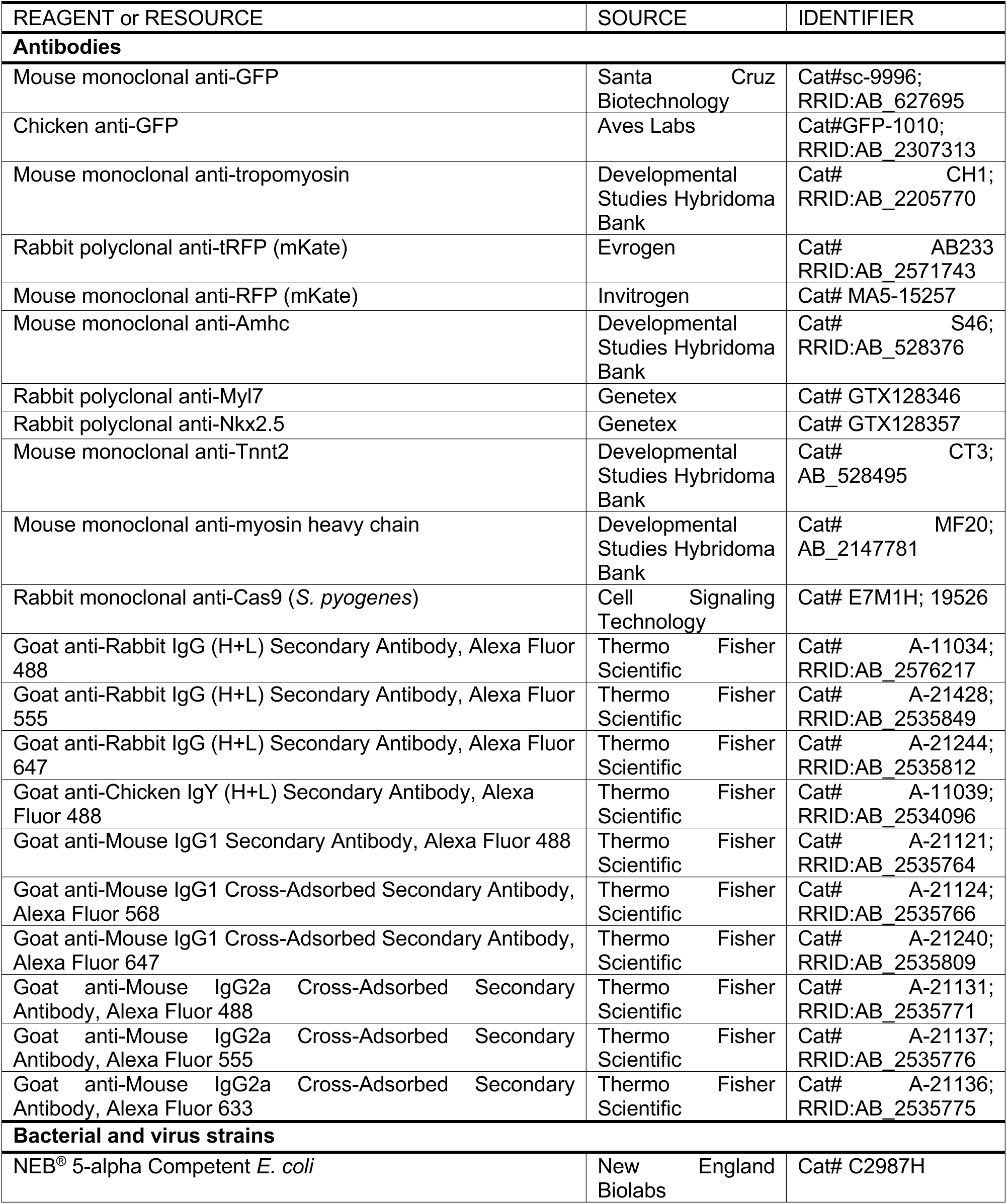
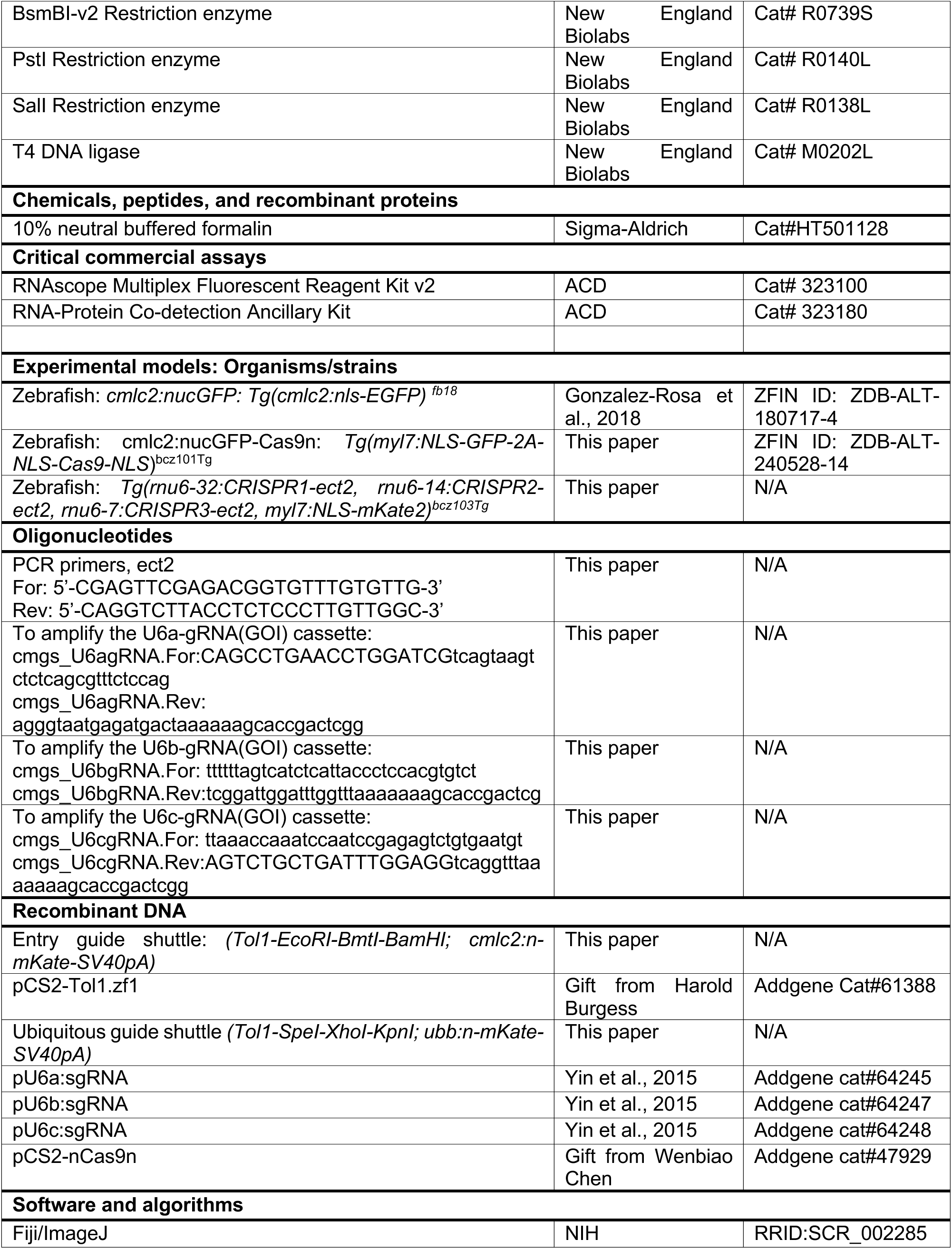
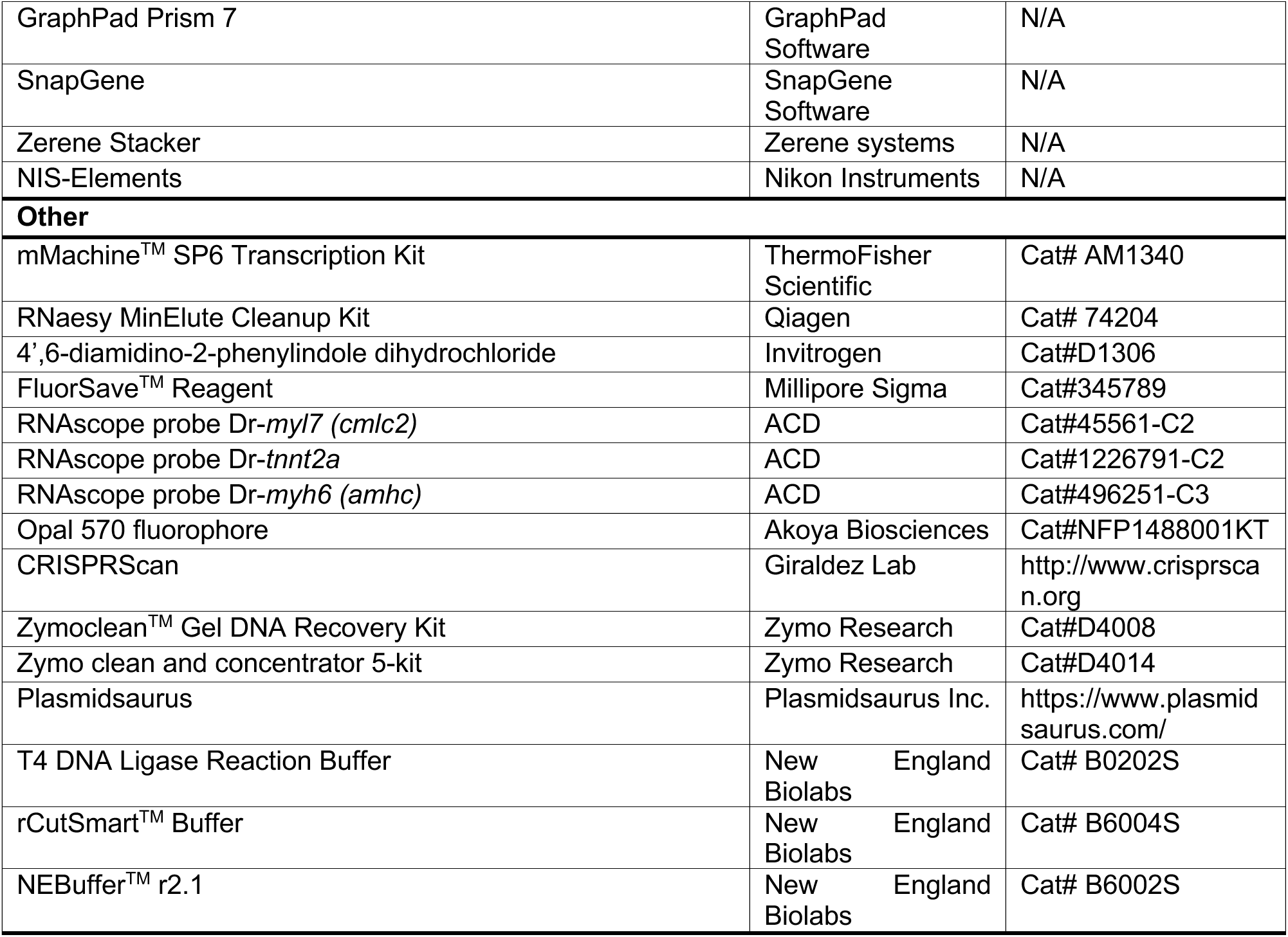

### Lead Contact

Further information and requests for resources or reagents, including fish lines, should be directed to and will be fulfilled by the lead contact, Dr. Juan Manuel González-Rosa.

### Zebrafish

Zebrafish embryos, larvae, and adults were produced, grown, and maintained according to standard protocols approved by the Institutional Animal Care and Use Committees of Massachusetts General Hospital and Boston College. Ethical approval was obtained from the Institutional Animal Care and Use Committees of Massachusetts General Hospital and Boston College. For experiments with adult zebrafish, animals ranging in age from 3 to 18 months were used. Approximately equal sex ratios were used for experiments. Adult density was maintained at 3-4 fish·l^-1^ for all experiments in Aquarius racks and fed twice daily. Water temperature was maintained at 28 °C. Published strains used in this study include wild-type AB and *Tg(cmlc2:nGFP)^fb^*^1827^ (ref. ^27^). Details of the construction of the new lines generated in this study are described below. At least three independent founders of each line were isolated and tested to confirm the described expression patterns and phenotypes. All the constructs generated in this study for transgenesis are available to the community.

### Construction of the cardiodeleter (cmlc2:nGFP-P2A-nCas9n) strain

To generate the *cmlc2:nGFP-nCas9n* transgenic line, a construct containing the following DNA elements was assembled by Gibson cloning: (1) a 0.9-kb *cmlc2* promoter to drive specific expression in cardiomyocytes; (2) a bicistronic *nlsGFP-P2A-nCas9n*; and (3) an SV40 polyadenylation signal. The nCas9n sequence was subcloned from pCS2-nCas9n (Addgene #47929, a gift from Wenbiao Chen^26^). The construct was flanked with Tol2 sites to facilitate transgenesis. In this line, all cardiomyocytes express a nuclear version of GFP and a nuclear localized, zebrafish codon optimized form of Cas9 from *Streptococcus pyogenes*). The official name of this line is *Tg(myl7:NLS-EGFP-2A-NLS-Cas9-NLS)^bcz^*^101^*^Tg^*.

### Construction of the *ect2 guide shuttle* strain

To generate the *ect2 guide shuttle* transgenic line, a construct containing the following DNA elements was assembled by Gibson cloning: (1) a U6a:sgRNA(*ect2* A) cassette (see details below for details on the cloning of the sgRNAs in the U6 plasmids); (2) a U6b:sgRNA(*ect2* B) cassette; (3) a U6c:sgRNA(*ect2* B) cassette; (4) a 0.9-kb *cmlc2* promoter to drive specific expression in cardiomyocytes; (5) the CDS of a nuclear-directed version of the red fluorescent protein mKate2; and (6) the polyadenylation signal from the ocean pout antifreeze protein (*afp*)^33^. Two versions of this construct were generated containing *Tol2* or *Tol1* sites to facilitate transgenesis. The official name of this line is *Tg(rnu6-32:CRISPR1-ect2, rnu6-14:CRISPR2-ect2, rnu6-7:CRISPR3-ect2, myl7:NLS-mKate2)^bcz^*^103^*^Tg^*.

### Construction of an empty, entry *guide shuttle* including cardiomyocyte labeling (*Tol1-EcoRI-BmtI-BamHI; cmlc2:n-mKate-SV40pA*)

To facilitate assembly of *guide shuttles* with a cardiomyocyte-specific reporter, a construct containing the following DNA elements was assembled by HiFi cloning (NEB): (1) a vector backbone containing Tol1 sites to facilitate transgenesis; (2) a ssDNA Ultramer oligonucleotide (IDT) containing *EcoRI*, *BmtI*, and *BamHI* restriction sequences; (3) 0.9-kb *cmlc2* promoter to drive specific expression in cardiomyocytes; (4) the CDS of a nuclear-directed version of the red fluorescent protein mKate; and (5) the polyadenylation signal from the ocean pout antifreeze protein (*afp*). This plasmid can be linearized by *EcoRI-HF*, *BmtI-HF*, and *BamHI-HF* digestion, and used as a backbone for subsequent cloning of the U6-sgRNA cassettes using Gibson Assembly (see details in the Advanced Protocol). Although the use of BmtI is optional, adding this enzyme minimizes the background from non-digested plasmids.

### Generation of cardiomyocyte-specific crispants

Embryos carrying the *cardiodeleter* transgene were injected at the one-cell stage with a 1 nL mix containing *guide shuttle* DNA and *Tol1* mRNA, both at 25 ng/μl. Injected animals were screened at 72 hpf to select embryos showing expression of mKate in cardiomyocytes, and then fixed at 4, 5 dpf or raised to adulthood.

For Tol1 mRNA synthesis, pCS2-Tol1.zf1 (Addgene #61388, a gift from Harold Burgess) was linearized with *NotI*, purified, and used from in vitro transcription using the mMessage mMachine SP6 Kit (ThermoFisher Scientific, AM1340). mRNA was purified using the RNeasy MinElute Cleanup Kit (Qiagen), and single-use aliquots were stored at −80 °C.

### Whole mount immunofluorescence and Imaging of Zebrafish Hearts

Embryos were fixed overnight in 4% paraformaldehyde (PFA) in PBS at 4 °C. Fixed embryos were bleached and permeabilized as described^51^, blocked in 5% bovine serum albumin and 5% goat serum in PBS containing 0.1% Tween-20 (PBST), and immunostained with primary antibodies diluted in blocking solution overnight at 4 °C. Primary antibodies used in this study include: anti-Amhc (S46, Developmental Studies Hybridoma Bank, 1:200), anti-GFP (clone B-2, Santa Cruz Biotechnology, 1:200, or AVES, 1:1,000), anti-mKate2 (tRFP, Evrogen, AB233; 1:1,000; or MA5-15257, Invitrogen, 1:1,1000), anti-Myl7 (Cmlc2, Genetex GTX128346, 1:500), anti-myosin heavy chain (MF20, DSHB, 1:100), anti-nkx2.5 (ref. ^48^, GTX128357, GeneTex, 1:500), anti-Tnnt2a (CT3, DSHB, 1:200), and anti-Tropomyosin (clone CH1, DSHB; 1:100). Alexa conjugated secondary antibodies (Life Technologies, 1:500) were used to detect primary antibody signals. Nuclei were counterstained with DAPI (Invitrogen). For imaging, hearts were then manually dissected and mounted in 0.9% low-melting agarose on glass-bottom dishes and imaged on either a Zeiss LSM900 or a Leica Stellaris 5 confocal microscope, both equipped with a 40x long working distance water objective. Images were processed and analyzed using Fiji/ImageJ^52^.

### Histological Analysis and Imaging

Adult zebrafish were euthanized by immersion in 0.16% tricaine (Sigma) and hearts dissected as described^53^. Freshly dissected adult zebrafish hearts were imaged using a SMZ18 fluorescence stereomicroscope (Nikon) equipped with the pE-300 white illumination technology (Cool LED) and a DS-Ri2 camera (Nikon). Multiple images capturing fluorescence or transmitted light were taken from each heart using the NIS-Elements Basic Research Package and then stacked using Zerene Stacker (Zerene Systems). Samples were fixed overnight at 4 °C in 4% paraformaldehyde (PFA) in PBS, included in paraffin and sectioned following conventional histological procedures. Immunofluorescence in paraffin sections were performed as described^54,55^. Primary antibodies used in this study include: anti-Amhc (S46, Developmental Studies Hybridoma Bank, 1:200), anti-Cas9 *S. pyogenes* (#19526, Cell Signaling Technology), anti-GFP (clone B-2, Santa Cruz Biotechnology, 1:200, or AVES, 1:1,000), anti-mKate2 (tRFP, Evrogen, AB233; 1:1,000; or MA5-15257, Invitrogen, 1:1,1000), anti-Myl7 (Cmlc2, Genetex GTX128346, 1:500), anti-myosin heavy chain (MF20, DSHB, 1:100), anti-nkx2.5 (ref. ^48^, GTX128357, GeneTex, 1:500), anti-Tnnt2a (CT3, DSHB, 1:200), and anti-Tropomyosin (clone CH1, DSHB; 1:100). Alexa conjugated secondary antibodies (Life Technologies, 1:500) were used to detect primary antibody signals. Nuclei were counterstained with DAPI (Invitrogen), and slides were mounted in FluorSave (Millipore). A Zeiss LSM900 confocal microscope was used to image immunostained sections.

Acid fuchsin-orange G (AFOG) stain was used to detect fibrotic tissue, as described^56^. Muscle, fibrin/cell debris, and collagen were stained brown, orange/red, and blue, respectively. Imaging was performed on a DM6 Leica Scope with a motorized stage and multi-image stitching was performed using the LAS X Navigator Software, or an Evident Slide Scanner.

### Combined RNA *in situ* hybridization and immunofluorescence

Protein-gene transcript co-detection on 7 μm paraffin heart sections was performed by RNAscope^TM^ (Advanced Cell Diagnostic, ACD). ACD designed gene probes targeting *cmlc2* (*myl7), tnnt2a, and amhc (myh6)* gene transcripts. The Multiplex Fluorescent Reagent Kit v2 was used for this purpose following manufacturer’s instructions. Briefly, sections were baked and deparaffinized and then treated with hydrogen peroxide prior to target retrieval. After target retrieval, sections were photobleached for 40 min in a 3% hydrogen peroxide, 20mM NaOH, PBS solution. Primary antibodies, including anti-Amhc (S46, Developmental Studies Hybridoma Bank, 1:200), anti-mKate2 (tRFP, Evrogen, AB233; 1:1,000; or MA5-15257, Invitrogen, 1:1,1000), anti-Myl7 (Cmlc2, Genetex GTX128346, 1:500), and anti-Tnnt2a (CT3, DSHB, 1:200), were incubated overnight at 4°C on tissue sections. Sections were then fixed in 10% Neutral Buffered Formalin for 30 min followed by Protease Plus treatment (15 min, 40 °C). Probes were hybridized for 2 hours at 40 °C and revealed using Opal 570 fluorophore from Akoya Biosciences (N FP1488001KT). Secondary antibodies incubation was performed at room temperature (RT) for 1 hour using Alexa conjugated secondary antibodies (Life Technologies, 1:500). Nuclei were counterstained with DAPI and slides were mounted in FluorSave (Millipore). Images were obtained in a Zeiss LSM 880 confocal microscope.

### Quantification of the efficiency of gene disruption mediated by Cas9

#### Tnnt2, Cmlc2, and Amhc

The frequency of biallelic deletion was estimated in a per-cardiomyocyte basis using the complete loss of the corresponding protein as a proxy. Stained embryos were imaged in a confocal microscope as described above, and z-stacks comprising most of the embryonic heart were collected for each animal. For Tnnt2a and Cmlc2, the percentages or mKate+ cardiomyocytes (i.e., cells that incorporated the guide shuttle) that displayed complete loss of the corresponding protein in the atrium and ventricle was quantified per animal. Because Amhc is only expressed in the atrium, only atrial mKate+ cardiomyocytes were quantified.

#### Ect2

Isolated hearts and fins from juvenile zebrafish were lysed by boiling in alkaline buffer, followed by neutralization with Tris-HCl. An amplicon around the first target of ect2 (gA) was produced using Phusion polymerase and primers 5’-CGAGTTCGAGACGGTGTTTGTGTTG-3’ and 5’-CAGGTCTTACCTCTCCCTTGTTGGC-3’, which was then subject to Next-Generation CRISPR Sequencing service at the MGH Center for Computational and Integrative Biology DNA Core. Primers designed for identifying big deletions can be found in Supplementary table X. Amplicons were subject to Sanger sequencing to confirm the elimination of the intervening sequence.

### Quantification and Statistical Analysis

Sample sizes were chosen based on previous publications and are indicated in each figure or figure legend. No animal or sample was excluded from the analysis unless the animal died during the procedure. All statistical values are displayed as mean ± standard deviation. Sample sizes, statistical test and *P* values are indicated in the figures or figure legends. Data distribution was determined before using parametric or non-parametric statistical test. Statistical significance was assigned at *P* < 0.05. All statistical tests were performed using Prism 9 software.

## SUPLEMENTARY FIGURES

**Figure S1 (related to Fig. 1).**
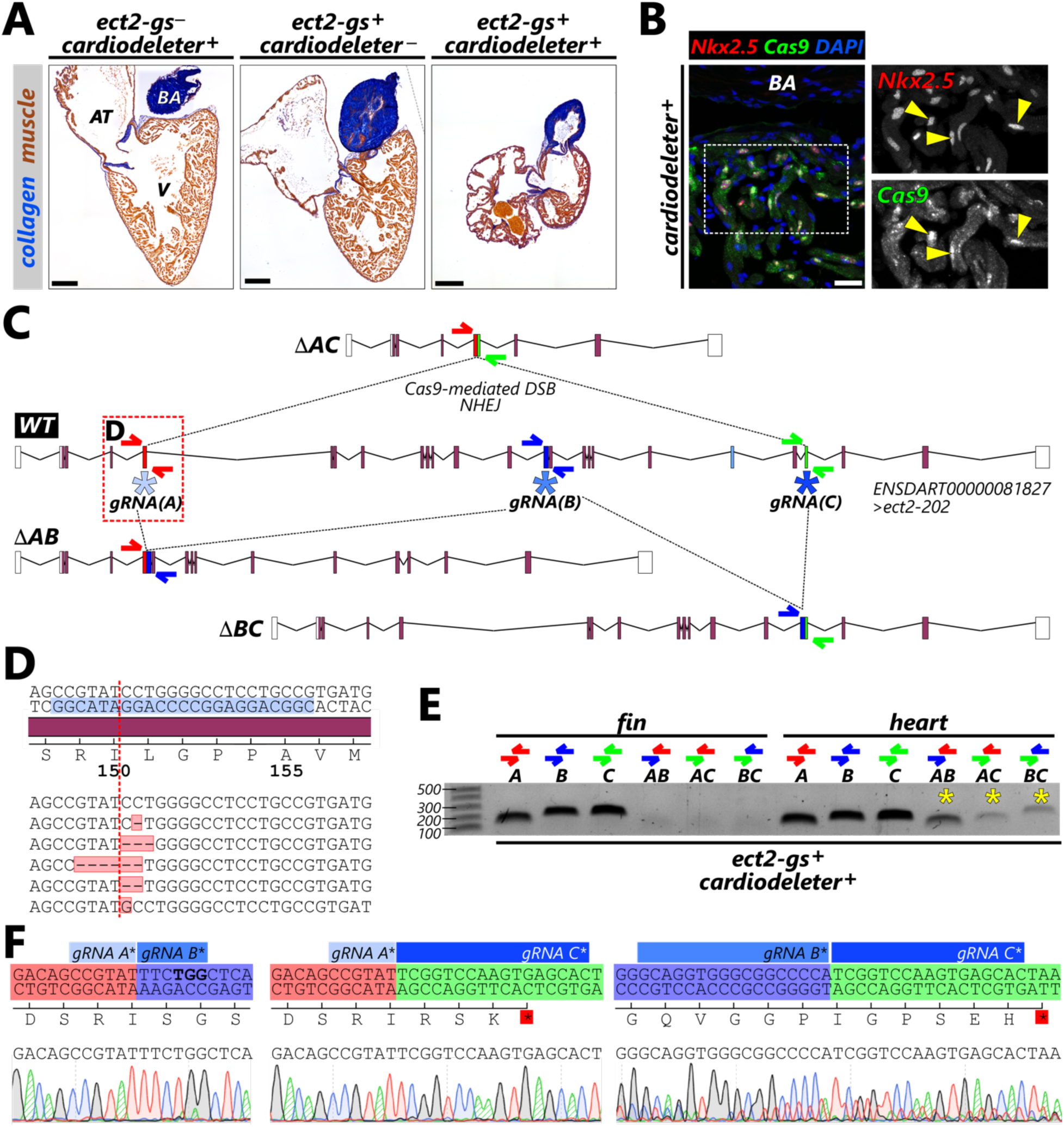
The *cardiodeleter* transgene disrupts the *ect2* locus when combined with a stable *ect2*-guide shuttle. **(A)** Heart sections from animals of the indicated genotypes stained using the AFOG technique to identify collagen (blue) and myocardium (orange/brown). **(B)** Confocal section of a zebrafish heart carrying the cardiodeleter transgene stained with antibodies to identify cardiomyocyte nuclei (Nkx2.5, red) and Cas9 (green). Nuclei are counterstained with the DNA dye DAPI (blue). Yellow arrowheads indicate Nkx2.5^+^; Cas9^+^ cardiomyocyte nuclei. **(C)** Representation of the wild-type (WT) *ect2* locus in the zebrafish genome and predicted *ect2* mutant alleles (ΔAB, ΔAC, and ΔBC) resulting from large deletions of the regions in between target sites. Asterisks indicate the location of the selected gRNA targets. The exons targeted by the *ect2*-gRNA-A,-B and-C appear red, blue, and green, respectively. Primers surrounding each of these targets are identified with the corresponding colors. In the Δ alleles, but not in the WT allele, the primers from different regions are only separated by a few hundred base pairs and can be used to detect the deletion event. **(D)** WT and mutant variants identified using NGS of an amplicon surrounding the target of gRNA-A. The gRNA(*ect2*)-A target sequence is highlighted in blue. Red line: predicted site of the Cas9-mediated double-strand break. **(E)** Representative PCRs using gDNA extracted from the fins or the heart of the same animals carrying the *cardiodeleter* and *ect2 guide shuttle* transgenes, using the indicated primer combination. Yellow asterisks indicate successful amplification of the Δ alleles. **(F)** Sanger sequencing results from the ΔAB, ΔAC, and ΔBC PCRs, demonstrating deletion of the intervening sequences in between the gRNA targets. AT, atrium; BA, bulbus arteriosus; V, ventricle. Scale bars: 200 μm (A) and 20 μm (B).

**Figure S2 (related to Figures 4 and 5).**
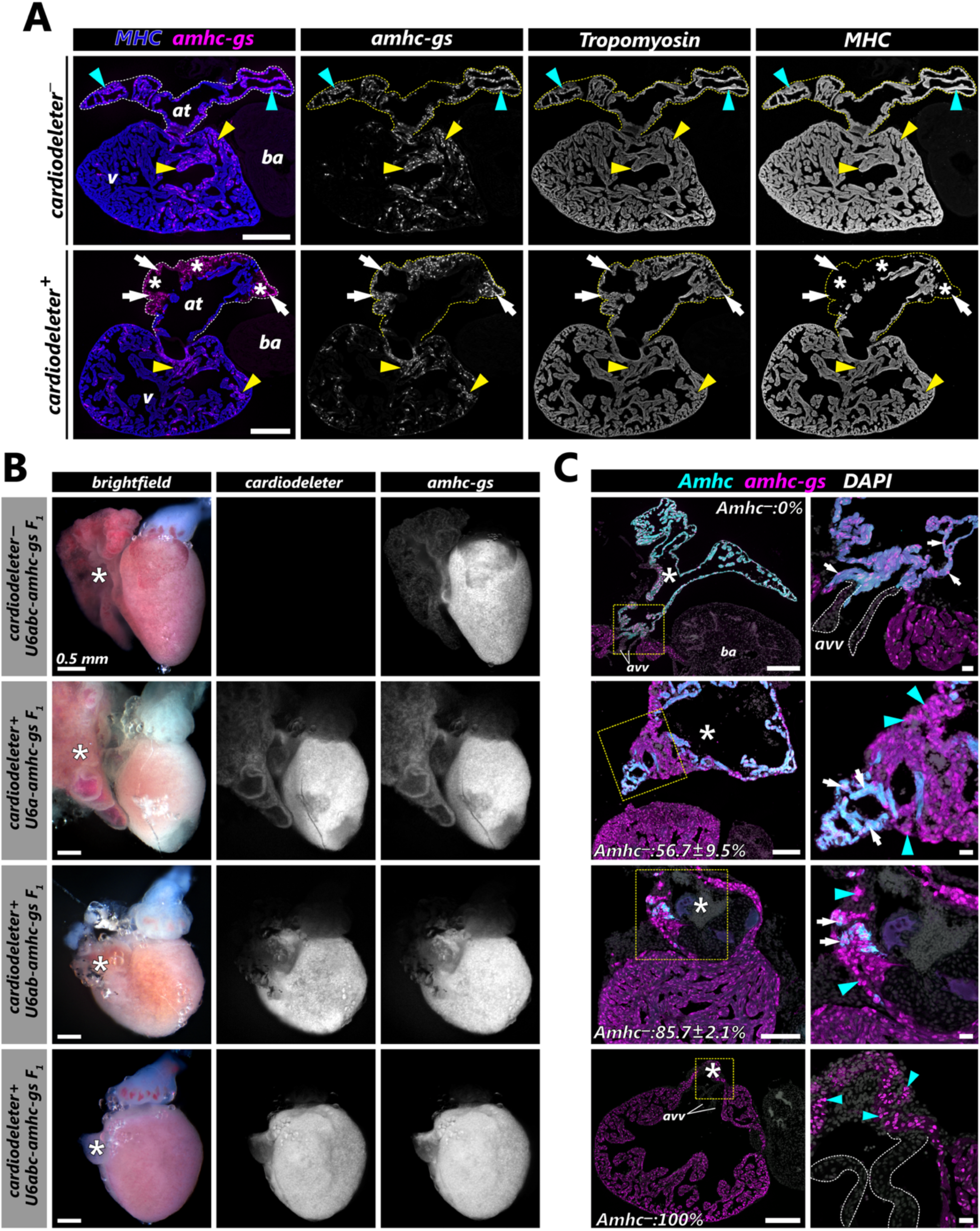
Characterization of *amhc* phenotypes induced upon Cas9-mediated mutagenesis. **(A)** Representative heart sections from 40 dpf wild-type (*cardiodeleter*-) and *cardiodeleter*+ animals injected with the *amhc*-gs. Sections were immunostained with antibodies to detect mKate expression (*amhc-gs*, magenta), tropomyosin, and myosin heavy chain (MHC, blue). Cyan arrowheads: *amhc-gs^+^*atrial cardiomyocytes expressing both tropomyosin and MHC. Yellow arrowheads: *amhc-gs^+^* ventricular cardiomyocytes expressing tropomyosin and MHC. White arrows: *amhc-gs^+^* atrial cardiomyocytes expressing tropomyosin but not MHC. Asterisks: *amhc-gs*^+^ clones in the atrium lacking MHC expression. **(B)** Representative bright field and fluorescence images of dissected hearts from the indicated cohorts. Asterisks: atrium. **(C)** Representative section from the hearts shown in B, immunostained with antibodies to detect mKate expression (*amhc-gs*, magenta), and Amhc (cyan). The average biallelic mutagenesis efficiency for each cohort is indicated in the corresponding panel. Magnifications of the dashed yellow boxed areas are shown on the right panels. White arrows: *amhc*-gs^+^ atrial cardiomyocytes expressing Amhc protein. Cyan arrowheads: *amhc*-gs+ atrial cardiomyocytes negative for Amhc protein expression; asterisks: atrium. at, atrium; v, ventricle; ba, bulbus arteriosus; avv, atrioventricular valve. Scale bars: 100 μm (A,C), 0.5 mm, (B).

## Notes

### Competing Interest Statement

The authors have declared no competing interest.

