## Extended Protocols - Generation of gene-specific guide shuttles. for "Optimization of methods for rapid and robust generation of cardiomyocyte-specific *crispants* in zebrafish using the *cardiodeleter* system"

**DETAILED PROTOCOL – Generation of gene-specific guide shuttles**

**Overview:** This protocol describes the methods to create a Tol1-based plasmid that carries the required sequences for producing three guide RNAs (gRNAs) targeting a gene of interest (GOI). The procedure involves three steps: (1) identifying potential gRNA, (2) cloning the gRNA sequences into U6-containing plasmids using oligonucleotide annealing; and (3) amplifying and assembling each U6-gRNA cassette into the ultimate guide shuttle. An example of how to generate an *ect2*-guide shuttle appears below in blue.

1. Selection of gRNAs targeting the gene of interest (GOI)
   1.1. Search your GOI in Ensembl.org. Press the "Show Transcript Table" button. Unless the goal is to target a specific splicing variant, identify the "Ensembl Canonical" transcript of your gene. Canonical transcripts are usually the most conserved and highly expressed, have the longest coding sequences, and are present in other repositories. Copy the Transcript ID (ENSDART followed by 11 digits).

   There are 5 annotated transcripts for *ect2* in the zebrafish genome, with ENSDART00000081827 identified as the canonical transcript.

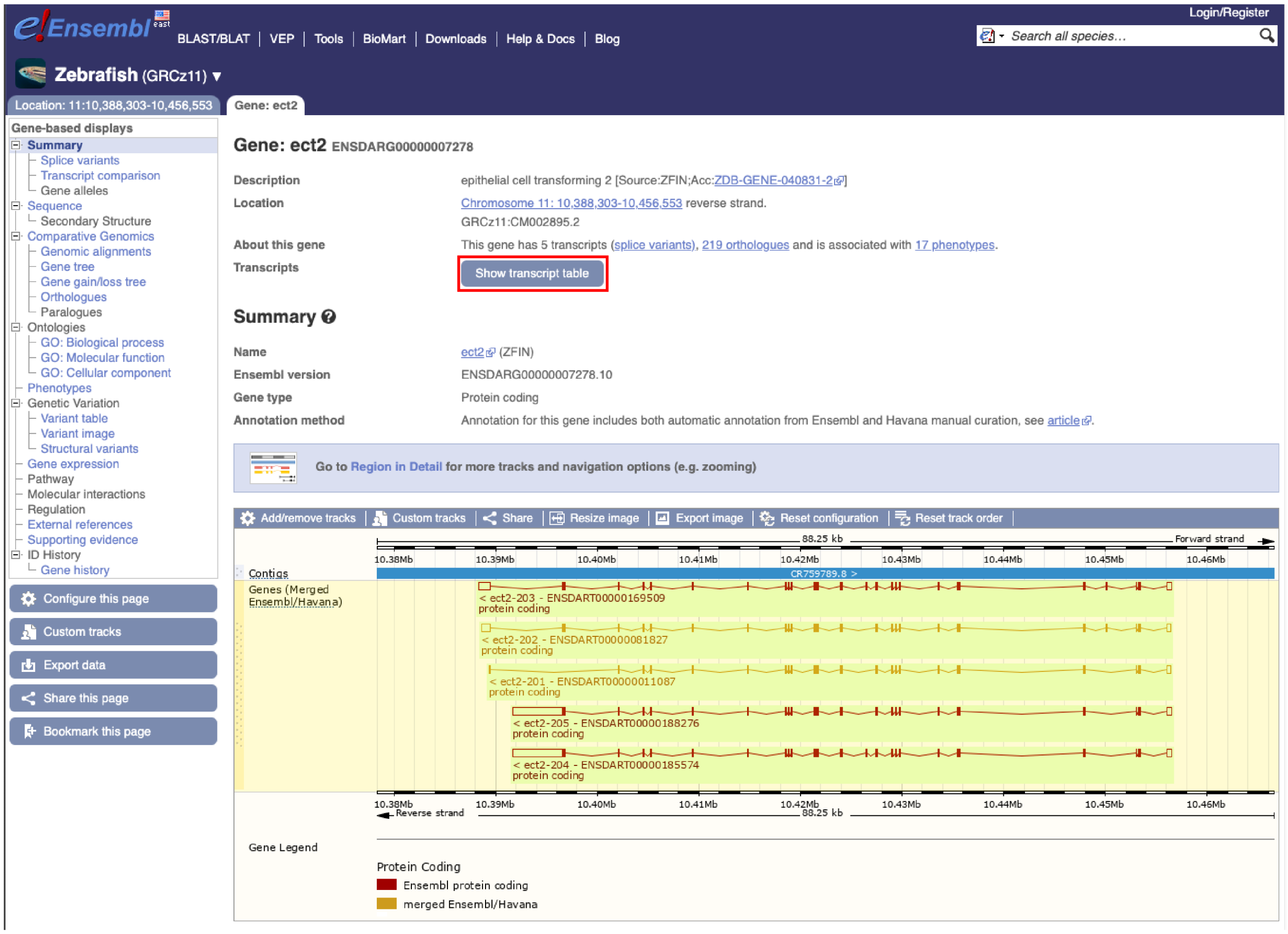


Show/hide columns (1 hidden)

| Transcript ID | Name | bp | Protein | Biotype | UniProt Match | Flags |
|---|---|---|---|---|---|---|
| ENSDART00000081827.4 | ect2-202 | 4186 | 976aa | Protein coding | F1QVS6 | Ensembl Canonical APPRIS ALT2 |
| ENSDART00000185574.1 | ect2-204 | 8427 | 980aa | Protein coding | A0A2R8Q1Z1 | APPRIS ALT2 |
| ENSDART00000188276.1 | ect2-205 | 8263 | 944aa | Protein coding | - | APPRIS ALT2 |
| ENSDART00000169509.3 | ect2-203 | 4793 | 1012aa | Protein coding | A0A8M2B2N3 | APPRIS P5 |
| ENSDART00000011087.6 | ect2-201 | 3054 | 878aa | Protein coding | Q6DRB9 | APPRIS ALT2 |

1.2. To identify target sites, access CRISPRScan (http://www.crisprscan.org/) and select the “By gene” option on the website's home page.

1.3. Paste the Transcript ID and select the *“Transcript”* option. We recommend keeping the *“In vitro T7 promoter”* option as both T7 and Pol III promoters, including U6, have a preference to start the transcription of small RNAs that begin with a G (ref. [39]). Maintain the rest of the default options and press *“Get sgRNAs”*. Several potential transcripts will be displayed. The transcript of interest will be highlighted in blue in the summary table at the bottom of the page.

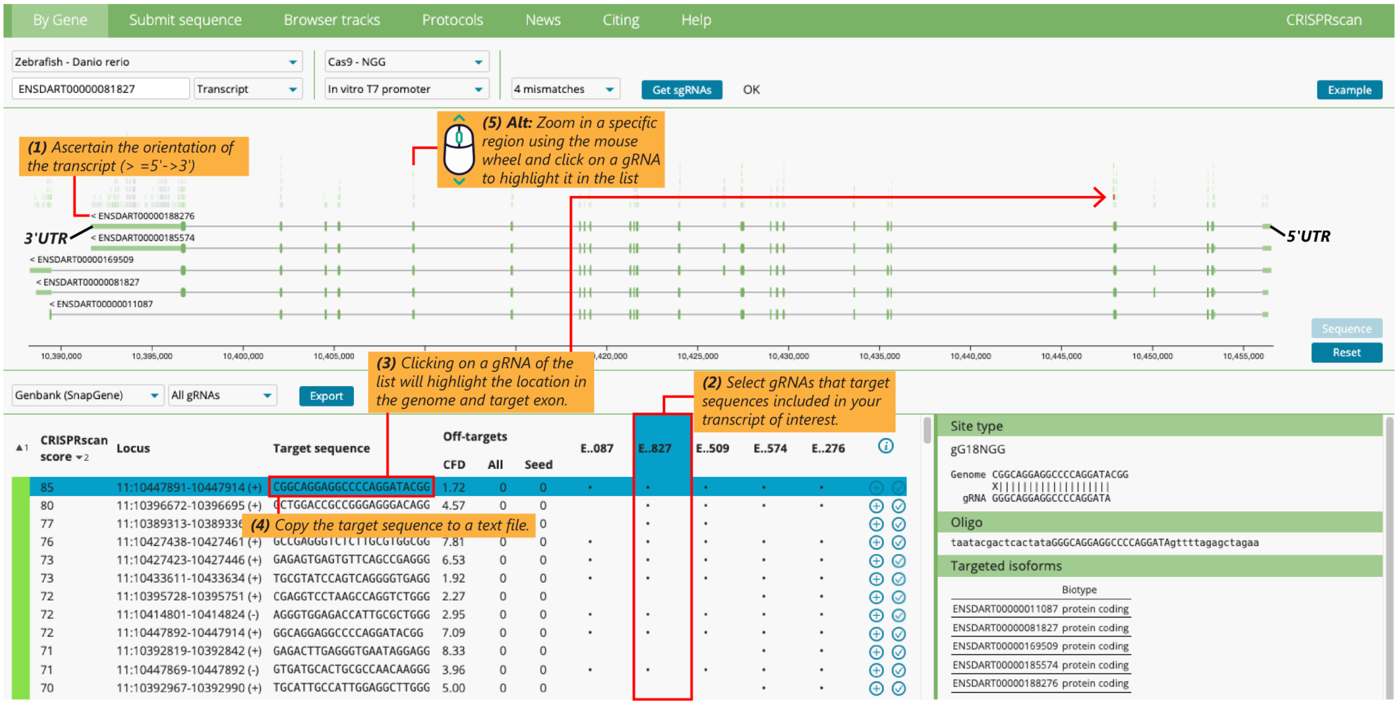


1.4. Select gRNAs with high CRISPRScan Score and no predicted off-targets (bright green). We recommend choosing three gRNAs targeting different exons across the entire gene.

For *ect2*, we selected three target sites (**PAM** sequenced in bold): GCAGGAGGCCCCAGGATA**CGG** (A, exon 5); AGGTGGGCGGCCCCATTC**TGG** (B, exon 14); and GCTCACTTGGACCGAGTC**AGG** (C, exon 23).

- Pay attention to the orientation of the gene. The symbol > in the Transcript ID number indicates the 5’→3’ orientation.
- By default, CRISPRScan will list the best guides (i.e., efficient and with no predicted off-targets) according to the CRISPRScore. To find out specifically what exon they are targeting, click on the sequences in the table at the bottom of the page. The selected gRNA will be highlighted in orange.
- To identify gRNAs in a specific location, use the mouse wheel to zoom in where desired and click on a gRNA to highlight it on the table.

1.5. Copy the selected **target sequences** in a text file. Each sequence should be 23 nt in the format 5’-NN-(18nt)-NGG-3’, where the red nucleotides correspond to the PAM.

For example, *ect2* target sequence A is:

5’-CGGCAGGAGGCCCCAGGATACGG-3’

1.6. **Oligonucleotide design.** The following steps provide a guide to design oligonucleotides that (1) contain the target sequence and (2), when annealed, will leave overhangs to facilitate their insertion into plasmids containing U6 promoters and the scaffold element of the sgRNA.

- Starting from the targeted sequence (from step 1.5), delete the green and red nucleotides:

18-nt target: 5’-GCAGGAGGCCCCAGGATA-3’

- To create the forward (FOR) primer, add in 5’: `ttcgg`

  `ect2-gA.FOR: 5’-`**`ttcgg`**`GCAGGAGGCCCCAGGATA-3’`

- To create the reverse (REV) primer:

  Obtain the reverse complement of the 18-nt target (for example, using Snapgene or https://reverse-complement.com/):

  `Reverse complement 18-nt target: 5’- TATCCTGGGGCCTCCTGC-3’`

  and then add the following modifications in 5’ and 3’:

  `ect2-gA.REV: 5’-`**`aaac`**`TATCCTGGGGCCTCCTGC`**`c`**` -3’`

When annealed, the primers should appear as follow:

```
ect2-gA.FOR:     5’-ttcggGCAGGAGGCCCCAGGATA-3’
ect2-gA.REV:         3’-cCGTCCTCCGGGGTCCTATcaaa-5’
```

1.7. Follow the same steps for each of the target sequences (three per gene). In total, you will have six primers, two for each gRNA that you will need to clone. Oligos can be ordered with no modifications from IDT or Sigma.

2. **Generation of pU6 constructs carrying a gRNA targeting the gene of interest.** The following steps will anneal the primers, creating overhangs to facilitate cloning. Once annealed, these oligonucleotides are integrated into the pU6 vectors using a one-pot restriction-ligation reaction. During the digestion, the pU6 plasmids are cut with three restriction enzymes. *BsmBI*, a Type II endonuclease, generates the sticky ends that are compatible with the annealed primers. *PstI* and *SalI* are added to this reaction to minimize the recovery of the uncut, original pU6-empty vector.

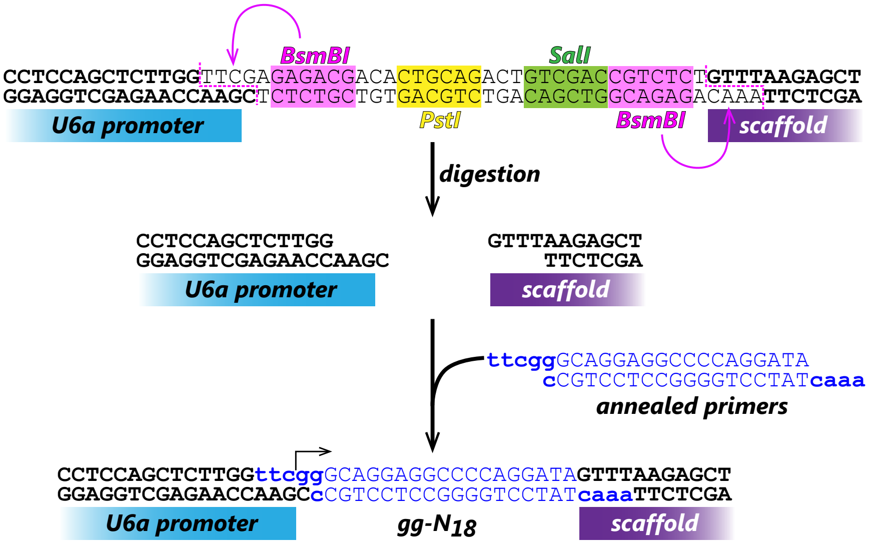


2.1. Centrifuge lyophilized primers 1 min, top speed (TS), room temperature (RT).

2.2. Reconstitute primers to 100 µM using milli-Q water (mQW). Resuspended primers can be stored at -20 °C for several years.

2.3. Prepare a buffered solution for primer annealing. The following table includes calculations for three reactions (R).

| | 1R | **3+1 = 4R** |
|---|---|---|
| mQW | 14 µl | 56 µl |
| NEB Buffer 2.1 10X | 2 µl | 8 µl |

2.4. Prepare the following annealing reaction mixes in PCR tubes:

| | **Annealing gA** | **Annealing gB** | **Annealing gC** |
|---|---|---|---|
| Buffer from step 2.3 | 16 µl | 16 µl | 16 µl |
| GOI-gA FOR 100 µM | 2 µl | – | – |
| GOI-gA REV 100 µM | 2 µl | – | – |
| GOI-gB FOR 100 µM | – | 2 µl | – |
| GOI-gB REV 100 µM | – | 2 µl | – |
| GOI-gC FOR 100 µM | – | – | 2 µl |
| GOI-gC REV 100 µM | – | – | 2 µl |

2.5. Transfer tubes to a thermocycler and run the following program. Annealed primers can be stored at -20 °C or used immediately in the following reaction.

| | |
|---|---|
| 95 °C | 5:00 |
| gradient to 50°C, 0.1 °C /s | - |
| 50 °C | 10:00 |
| 4 °C | ∞ |

2.6. Prepare the one-pot digestion-ligation master mix. Keep enzymes in a cooler and assemble the reaction on ice. Mix well.

| | 1R | **3+1 = 4R** |
|---|---|---|
| mQW | 5 µl | 20 µl |
| CutSmart Buffer NEB 10x* | 1 µl | 4 µl |
| T4 DNA ligase buffer 10x | 1 µl | 4 µl |
| T4 DNA ligase | 0.3 µl | 1.2 µl |
| *BsmBI v2* | 0.3 µl | 1.2 µl |
| *PstI* | 0.2 µl | 0.8 µl |
| *SalI* | 0.2 µl | 0.8 µl |

* Although *BsmBI v2* is predicted to have very little activity in this buffer, we have experimentally found that using CutSmart in this reaction produces significantly higher efficiency than other buffers.

2.7. Assemble the digestion-ligation reactions (on ice):

| | **pU6a-gA** | **pU6b-gB** | **pU6c-gC** |
|---|---|---|---|
| Digestion-ligation master mix (from step 2.6) | 8 µl | 8 µl | 8 µl |
| Annealed gA oligos (from step 2.4) | 1 µl | – | – |
| Annealed gB oligos (from step 2.4) | – | 1 µl | – |
| Annealed gC oligos (from step 2.4) | – | – | 1 µl |
| pU6a-gRNA (100 ng/ µl; Addgene #64245) | 1 µl | – | – |
| pU6b-gRNA (100 ng/ µl; Addgene #64247) | – | 1 µl | – |
| pU6c-gRNA (100 ng/ µl; Addgene #64248) | – | – | 1 µl |

2.8. Transfer tubes to a Thermocycler and start the following program.

| | | |
|---|---|---|
| 37 °C | 20:00 | x3 |
| 16 °C | 15:00 | |
| 37 °C | 10:00 | |
| 55 °C | 15:00 | |
| 80 °C | 15:00 | |
| 4 °C | ∞ | |

2.9. Using standard methods, perform transformation of TOP10 or NEB DH5α bacteria with 2 μl of the reaction. To reduce costs, a 50 μl vial of NEB DH5α bacteria can be split to perform up to 6 transformations.

2.10. Plate 100 μl of each transformation on prewarmed LB Spectinomycin (50 μg/ml) plates and incubate overnight at 37 °C.

2.11. Identify colonies carrying the correct insert using colony PCR.
PCR using M13.FOR + the GOI-gX.REV primers can be used to distinguish between bacteria transformed with the correct and the empty pU6 plasmid.

Example of the pU6a-ect2.gA showing annealing of the M13.FOR and ect2-gA.REV primers, which would produce a 482 bp amplicon.

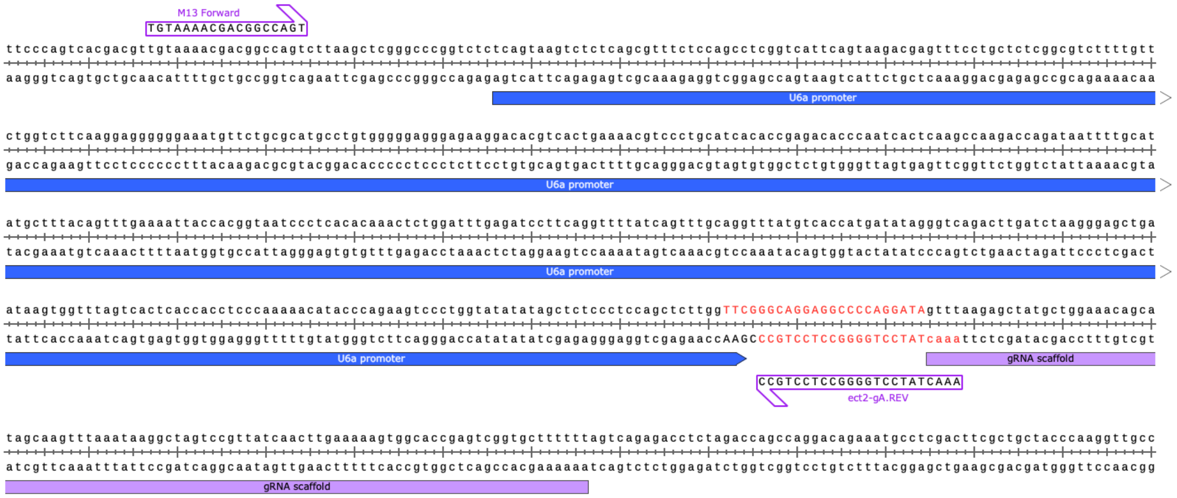


Original pU6a construct, containing the *BsmBI* cassette. No amplicon will be produced because the REV primer does not anneal in this construct.

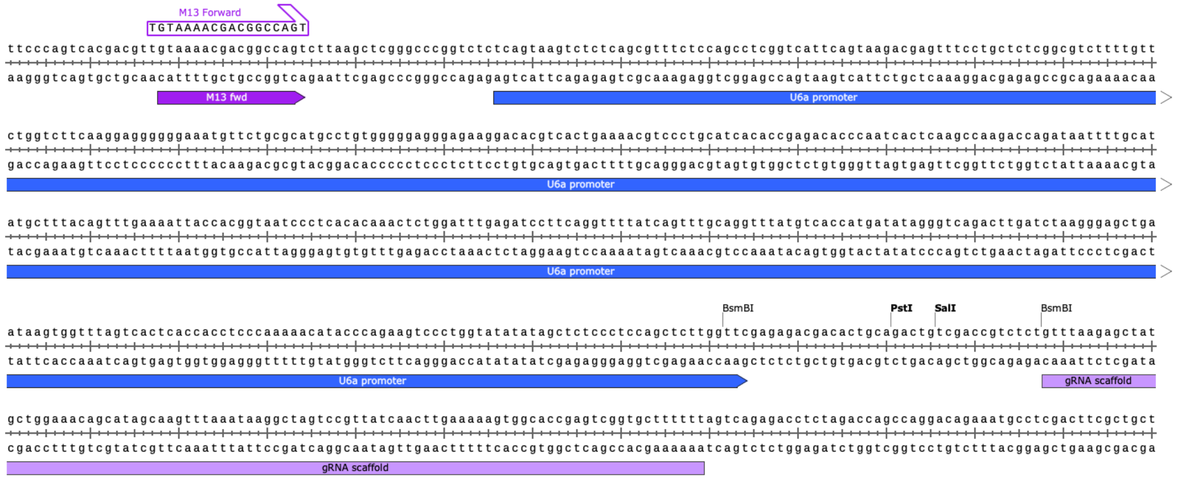


- Prepare a 10 mM solution of the REV primer for each guide.
- In a 1.5 ml tube, prepare the following mix (common for screening 4 colonies of each of the 3 plasmids):

| | 1R | **4x3 + 1 = 13R** |
|---|---|---|
| Platinum Master Mix 2x | 5 μl | 65 μl |
| M13 FOR primer | 0.4 μl | 5.2 μl |
| mQW | 4.2 μl | 54.6 μl |
| **TOTAL** | **9.6 μl** | **124.8 μl** |

- Prepare the following PCR mixes to test each construct:

| | for gA | for gB | for gC |
|---|---|---|---|
| Master Mix from step #2 | 41.2 µl | 41.2 µl l | 41.2 µl |
| GOI-gA.REV | 1.6 µl | – | – |
| GOI-gB.REV | – | 1.6 µl | – |
| GOI-gC.REV | – | – | 1.6 µl |
| **TOTAL** | **42.8 µl** | **42.8 µl** | **42.8 µl** |

- Prepare a set of PCR tubes (4 tubes per construct) and dispense 10 µl of PCR mix in each tube. Keep them on ice.
- Prepare another set of PCR tubes and add 20 µl of mQW to each tube. Working in sterility conditions and using 10 µl sterile tips, pick colonies of the corresponding plates, immerse the tip in the water tube, and shake gently to release some bacterial mass. Then, transfer the tip immediately to the corresponding PCR mix and shake it again. These tubes will serve as a replica of each colony. If the construct is correct, the solution of water and bacteria will be used to inoculate a miniculture.
- Transfer the PCR tubes to a thermocycler and use the following program:

| | | |
|---|---|---|
| 94 °C | 2:00 | |
| 94 °C | 0:15 | x34 |
| 60 °C | 0:15 | |
| 68 °C | 0:30 | |
| 68 °C | 3:00 | |
| 4 °C | ∞ | |

- Run PCR products in a 1% agarose gel. Colonies containing the correct plasmid will produce an amplicon of ~450 to ~480 bp.

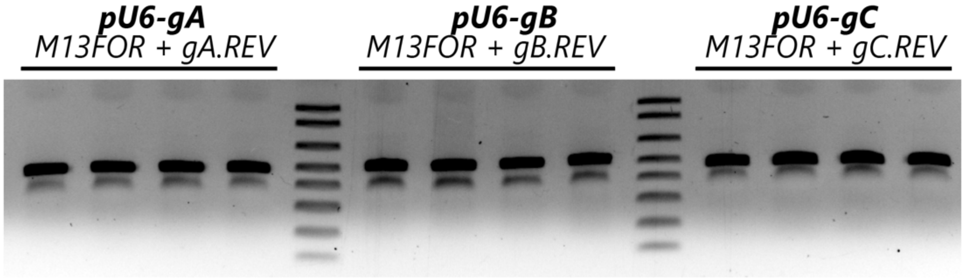


- Grow 2 correct clones per construct in 5 mL of LB + Spectinomycin 50 µg/ml, overnight at 37 °C.
- Create glycerol stocks by mixing ~200 µl of miniculture and ~200 µl of glycerol 50%. Store glycerol stocks at -80 °C.
- Extract DNA using standard miniprep procedures.
- Send one or two clones for Sanger sequencing using the M13FOR primer to confirm correct sequence of the insert.

3. **Subcloning the U6-gRNA cassettes into the *Tol1-cm:n-mKate* guide shuttle.** The goal of the following steps is to amplify the U6-gRNA cassettes and assemble them into a guide shuttle that includes a cardiomyocyte-specific transgenic reporter.

<u>*Vector linearization*</u>

3.1. Set up digestion of the empty *Tol1-cm:n-mKate* guide shuttle:

| | **1R** |
|---|---|
| DNA (*Tol1-cm:n-mKate* guide shuttle) | ~5 μg |
| CutSmart Buffer 10x | 5 μl |
| *EcoRI*-HF | 1 μl |
| *BmtI*-HF | 1 μl |
| *BamHI-HF* | 1 μl |
| mQW | up to 50 μl |
| **TOTAL** | **50 μl** |

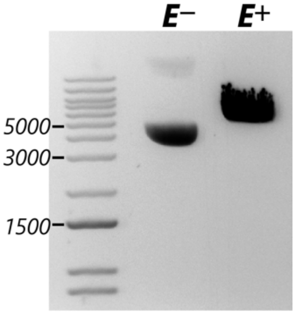


3.2. Incubate reaction for 3 hours at 37 ºC.
3.3. Run a small aliquot of the digestion (~5 μl) and a sample of the undigested plasmid in a 1% agarose gel to confirm complete linearization.
3.4. Purify the rest of the linearized DNA using the Zymo DNA Clean and Concentrate kit. Elute in mQW.
3.5. Determine DNA concentration and dilute the vector to a final concentration of 50 ng/μl.

*<u>Preparation of the gene-specific U6-gRNA fragments</u>*

3.6. Resuspend the U6x-gRNA specific primers and prepare working solutions at 4 μM.
To amplify the U6a-gRNA(GOI) cassette:
cmgs_**U6agRNA**.FOR: CAGCCTGAACCTGGATCGtcagtaagtctctcagcgtttctccag
cmgs_**U6agRNA**.REV: agggtaatgagatgactaaaaaagcaccgactcgg

To amplify the U6b-gRNA(GOI) cassette:
cmgs_**U6bgRNA**.FOR: ttttttagtcatctcattaccctccacgtgtct
cmgs_**U6bgRNA**.REV: tcggattggatttggtttaaaaaaagcaccgactcg

To amplify the U6c-gRNA(GOI) cassette:
cmgs_**U6cgRNA**.FOR: ttaaaccaaatccaatccgagagtctgtgaatgt
cmgs_**U6cgRNA**.REV: AGTCTGCTGATTTGGAGGtcaggtttaaaaaaagcaccgactcgg

3.7. Set up the following PCR reactions:

| | **U6a-gRNA cassette** | **U6b-gRNA cassette** | **U6c-gRNA cassette** |
|---|---|---|---|
| Phusion 2X GC Mix | 25 μl | 25 μl | 25 μl |
| mQW | 19 μl | 19 μl | 19 μl |
| cmgs_**U6a**gRNA.FOR (4 μM) | 2.5 μl | – | – |
| cmgs_**U6a**gRNA.REV (4 μM) | 2.5 μl | – | – |
| cmgs_**U6b**gRNA.FOR (4 μM) | | 2.5 μl | – |
| cmgs_**U6b**gRNA.FOR (4 μM) | | 2.5 μl | |
| cmgs_**U6c**gRNA.FOR (4 μM) | – | – | 2.5 μl |
| cmgs_**U6c**gRNA.FOR (4 μM) | – | – | 2.5 μl |
| **pU6a**(GOI-gRNA) (5 ng/μl) | 1 μl | – | – |
| **pU6b**(GOI-gRNA) (5 ng/μl) | – | 1 μl | – |
| **pU6c**(GOI-gRNA) (5 ng/μl) | – | – | 1 μl |

3.7. Transfer the PCR tubes to a thermocycler and run the following program:

| | | |
|---|---|---|
| 98 °C | 0:30 | |
| 98 °C | 0:10 | x34 |
| * °C | 0:30 | |
| 72 °C | 0:20 | |
| 72 °C | 5:00 | |
| 4 °C | ∞ | |

* Using the Phusion 2X GC Mix, use 65 ºC to amplify the U6a- and U6c-gRNA cassettes. Use 64 ºC to amplify the U6b-gRNA cassette.

3.8. Run PCR products in a 0.8-1% agarose gel.
3.9. Using a transilluminator, isolate the correct amplicons (~560 bp) from the gel and extract the DNA using the Zymo Gel DNA Recovery kit. Follow the manufacturer’s instruction and elute in mQW pre-heated at 42 ºC.
3.10. Determine the concentration of the eluted DNA and dilute each purified PCR to 20 ng/μl.

*Assembly of the ultimate guide shuttle*

3.11. Assemble the following HiFi Assembly (NEB) reaction:

| | **1R** |
|---|---|
| HiFi 2X Mix | 5 μl |
| Linearized *Tol1-cm:n-mKate* guide shuttle (50 ng/μl) | 1 μl |
| Amplified U6a-gRNA(GOI) cassette (20 ng/μl) | 1 μl |
| Amplified U6b-gRNA(GOI) cassette (20 ng/μl) | 1 μl |
| Amplified U6c-gRNA(GOI) cassette (20 ng/μl) | 1 μl |
| mQW | 1 μl |
| **TOTAL** | **10 μl** |

3.12. Incubate the HiFi Assembly reaction for 1 hour at 50 ºC.
3.13. Using standard methods, perform transformation of NEB DH5α bacteria with **exactly** 2 μl of the assembly.
3.14. Plate 100 μl of the transformation on pre-warmed LB-Ampicillin (100 μg/ml) plates and incubate overnight at 37 °C.
3.15. Pick 8 colonies and inoculate 5 mL of LB Ampicillin (100 μg/ml) minicultures. Grow overnight at 37 °C, 250 rpm.
3.16. Create glycerol stocks by mixing ~200 μl of miniculture and ~200 μl of glycerol 50%. Store glycerol stocks at -80 °C.
3.17. Extract DNA using standard miniprep procedures.
3.18. Set up restriction digestions to identify colonies carrying the correct construct.

| | **1R** | **8+1=9R** |
|---|---|---|
| Miniprep DNA | 500 ng -1μg | |
| NEB CutSmart 10x Buffer | 5 μl | 45 μl |
| *BsrGI*-HF | 1 μl | 9 μl |
| *HindIII*-HF | 1 μl | 9 μl |
| mQW | up to 50 μl | up to 450 μl |
| **TOTAL** | **50 μl** | **450 μl** |

3.19. Incubate digestions 1 h at 37 °C.

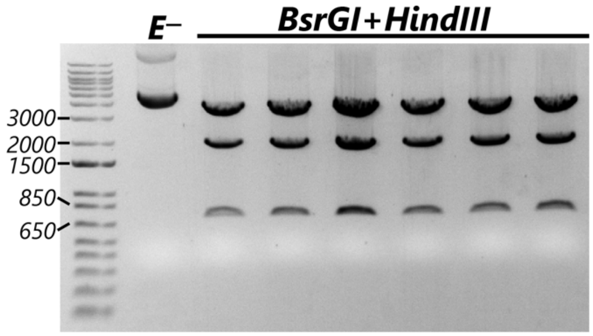


3.20. Run digestion reactions in a 1% agarose gel to verify restriction pattern. Include a non-digested (E-) sample to determine the approximate size of the coiled plasmid. Although subject to variations depending on the gRNA sequences, digestion of most final guide shuttle result in three bands of ~3.9, ~2, and ~0.7 kb.

3.21. Send one or two constructs with the correct pattern for whole plasmids sequencing (i.e., using Plasmidsaurus) to verify that all elements are correctly assembled and there are no undesired mutations.

3.22. Once the construct has been verified by sequencing, DNA for injection can be prepared from a maxiprep. Alternatively, if available, ~5 μg from the remaining miniprep can be cleaned up using the Zymo Clean and Concentrator kit.

**4. *Guide shuttle* injection and selection.**

4.1. Produce Tol1 mRNA using standard methods of in vitro transcription. Store 2 μl aliquots of mRNA at 250 ng/μl at -80 °C.

4.2. The day before the injection, set up crosses of WT or tissue-specific Cas9+ animals with a divider.

4.3. On the injection day, thaw an aliquot of Tol1 mRNA on ice and add the following reagents directly to that tube:

| | 1R |
|---|---|
| Tol1 mRNA (250 ng/μl) | 2 μl |
| Phenol red | 2 μl |
| KCl 2 M | 2 μl |
| Purified guide shuttle plasmid (250 ng/μl) | 2 μl |
| mQW | 12 μl |
| **TOTAL** | **20 μl** |

4.4. Load 3 μl of the injection mix in microinjection needle and inject into zebrafish embryos at the one-cell stage following standard methods.

**_Appendix I_: *In silico* assembly of the pU6-gRNA plasmids and guide shuttles** for sequencing and verification purposes.

**_Method 1_:** Using Snapgene to design and plan all cloning steps.

**5. Assembly of the pU6 constructs.**

5.1. Download the sequences of Addgene plasmids # 64245 (pU6a), 64247 (pU6b), and 64248 (pU6c). As a consensus, we clone gA in pU6a, gB in pU6b, and gC in pU6c. An annotated version of these constructs identifying each component, including the U6 promoter and the gRNA scaffold, can be found in the Supplemental Sequence file.

5.2. In Snapgene's *Actions* Menu, click on *Anneal Oligos*.

5.3. For each of the gene-gX primer pairs generated in section 1.6 (for example, ect2-gA.FOR and ect2-gA.REV), paste the sequence of the oligos that you will anneal, create a name (i.e., Annealed_ect2-gA), and select *Anneal*. Save the results and repeat this procedure to anneal the primer pairs from gB and gC.

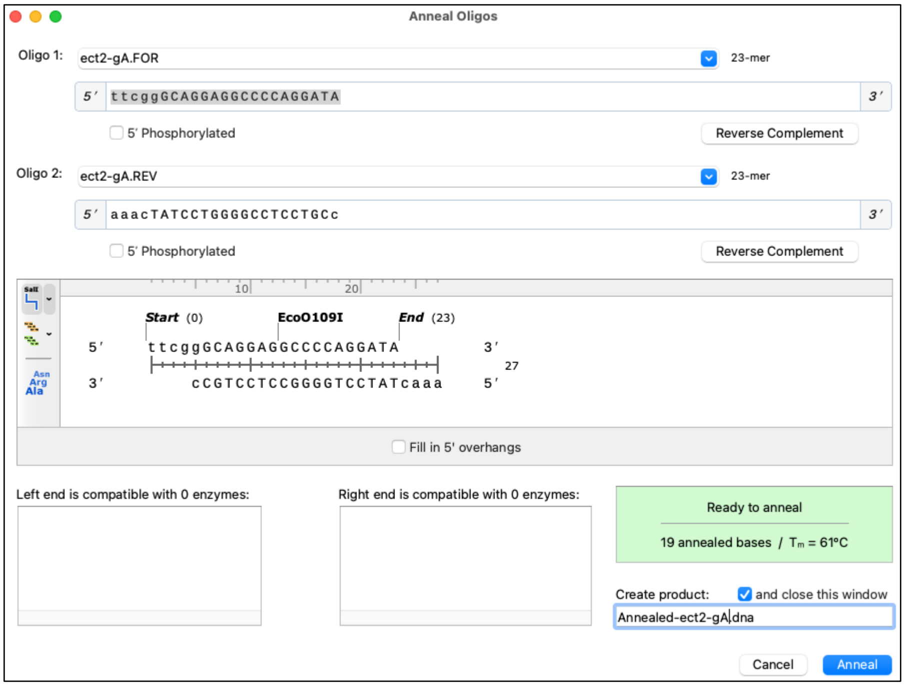


5.4. Using Snapgene, open the sequence of plasmid pU6a. In the *Actions* menu, select *Restriction and Insertion Cloning > Insert Fragment*.

5.5. In the _Vector_ Tab, select *BsmBI* as the restriction enzyme.

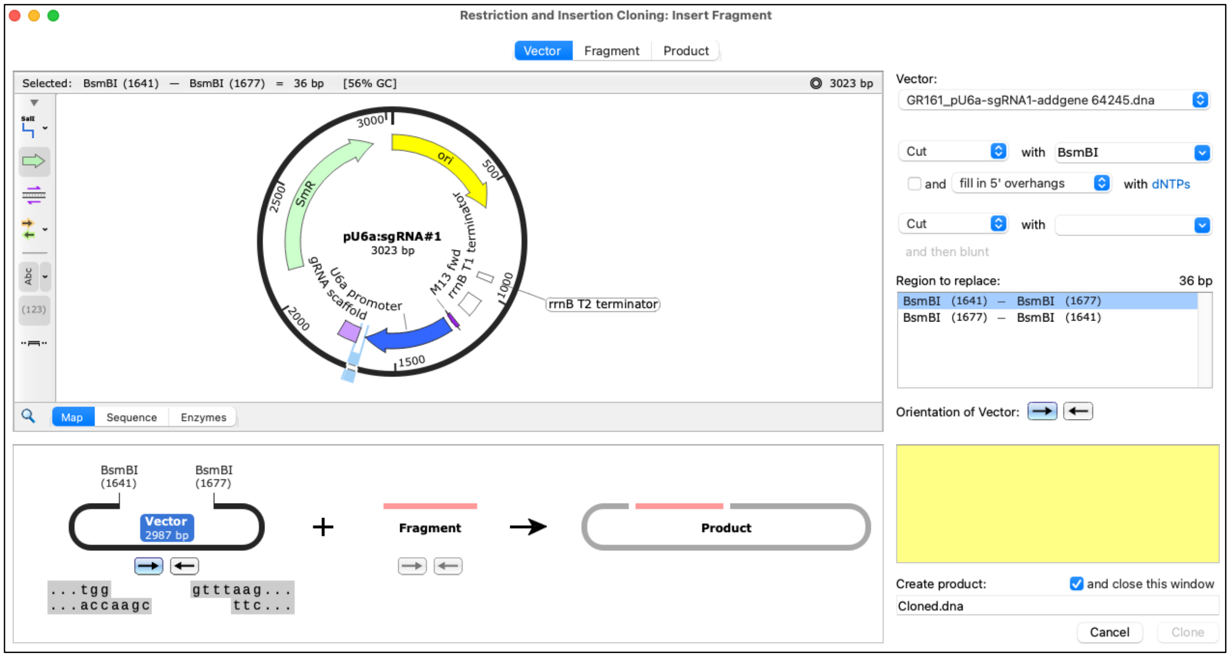


5.6. Move to the _Fragment_ Tab. Select the file containing the annealed primers under *Source of Fragment* dropdown menu. Write down a name for the construct and select *Clone*.

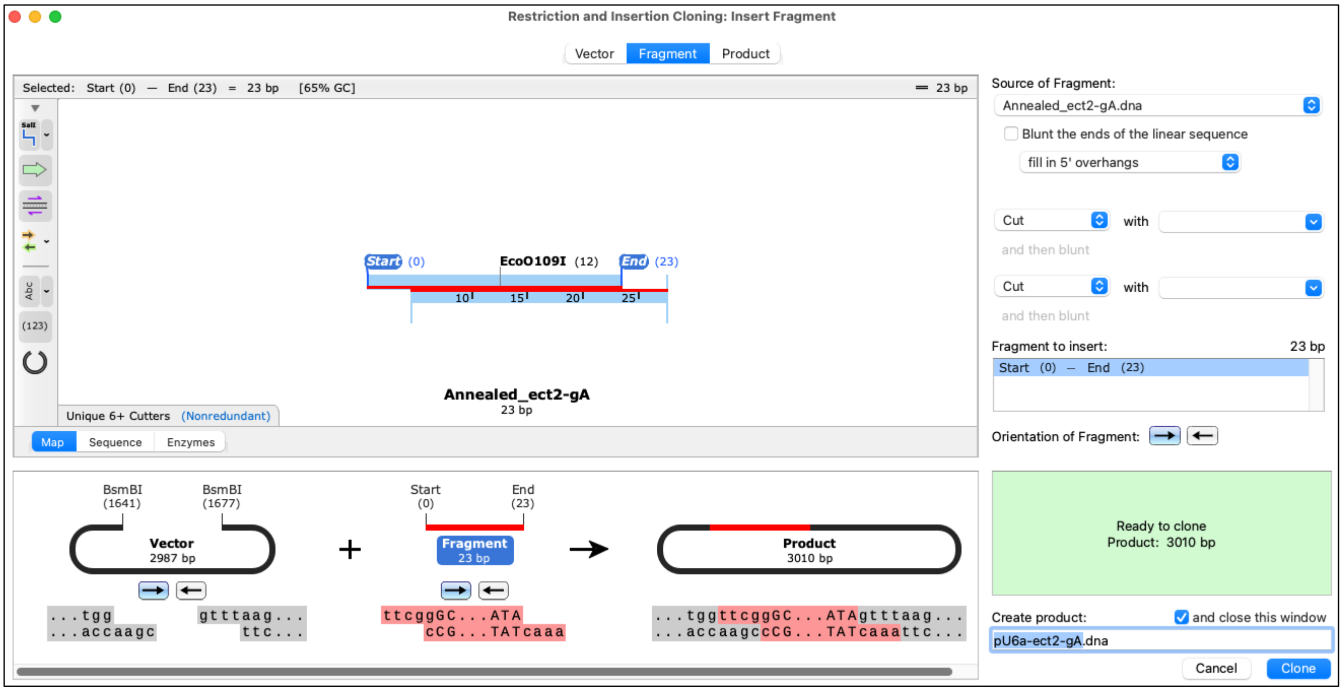


5.7. Repeat the same procedure to simulate the cloning of the annealed primers containing the gB and gC target sequence in pU6b and pU6c.

### 6. Assembly of the *Tol1-cm:n-mKate* guide shuttle.

<u>*Simulating the Vector Linearization*</u> (steps 3.1-3.5)

6.1. Open the .dna file containing the sequence of the empty *Tol1-cm:n-mKate guide shuttle* in Snapgene.

6.2. Activate the *Show Enzymes* option from the left panel, and select the *Unique 6+ Cutters* option.

6.3. Click on *EcoRI* and *BamHI* while pressing <u>Shift</u> in your keyboard.

6.4. In the *Actions* menu, select *Linearize*. This will produce a new file containing the result from the digestion, including the corresponding overhangs. Save this file.

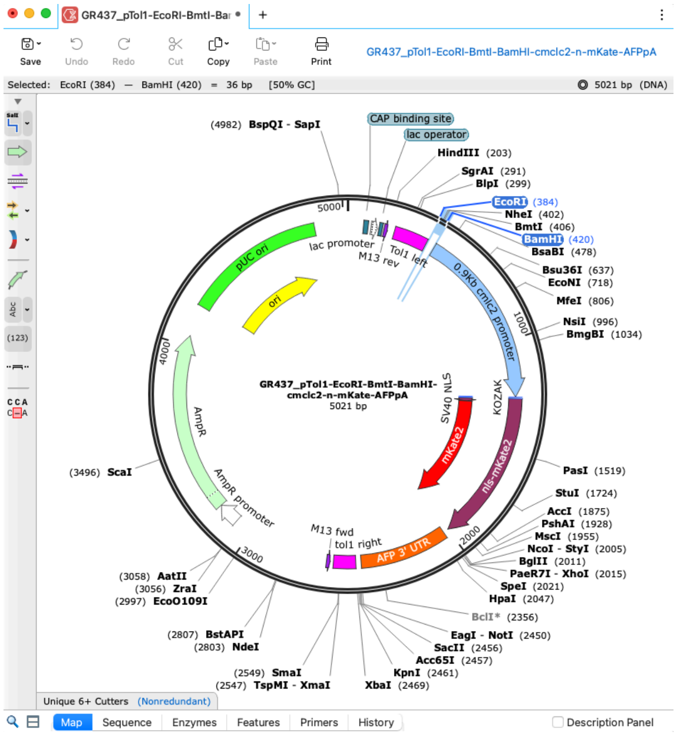

*Simulating the Preparation of the Gene-Specific U6-gRNA Fragments* (steps 3.6-3.10)

6.5. Open the sequence of your pU6a-GOI-gA in Snapgene.

6.6. In the *Actions* menu, select *PCR*.

6.7. Paste the sequences of the cmgs_U6agRNA.FOR and .REV. Add a new name for the U6a-GOI-gA amplicon and click on *PCR*.

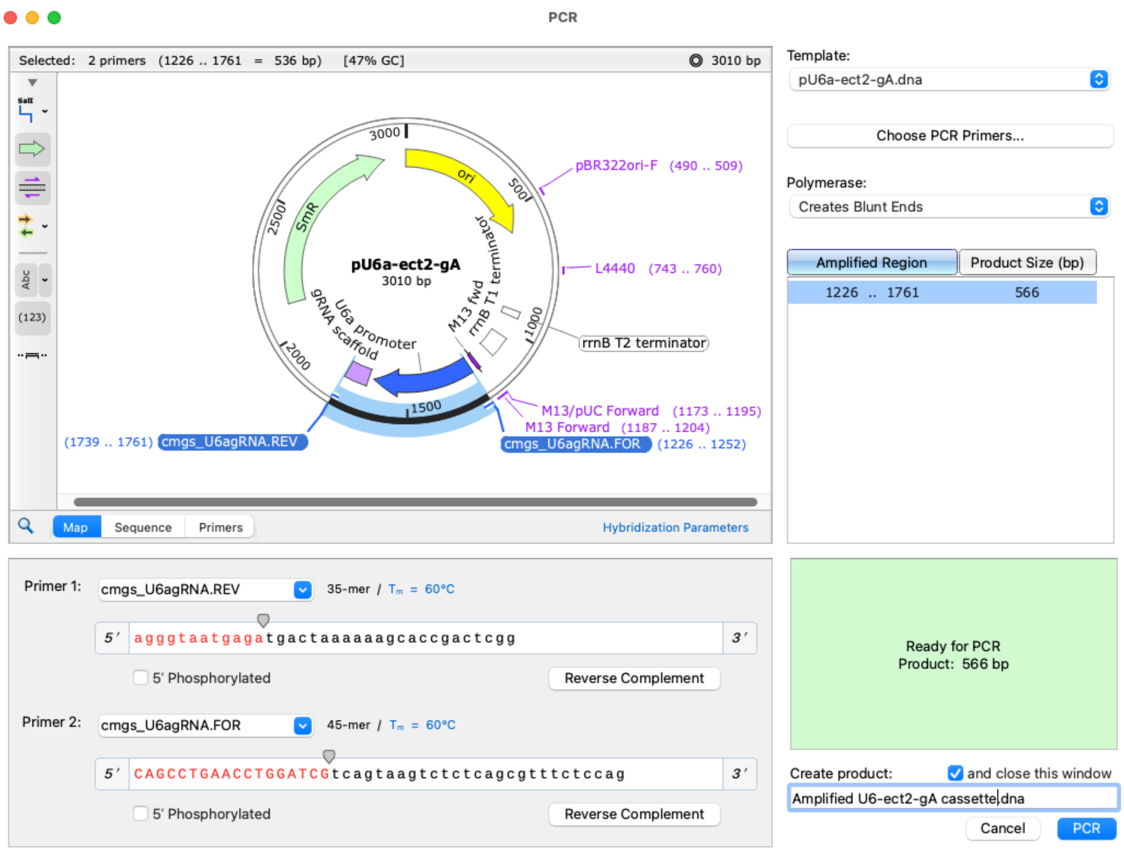


6.8. Repeat these steps using the corresponding templates and primers to obtain the U6-GOI-gB cassette and U6c-GOI-gC cassette.

*Simulating Assembly of the Guide Shuttle* (steps 3.11-3.12)

6.9. Open the linearized, empty guide shuttle file in Snapgene. In the *Actions* menu, select *NEBuilder® HiFi DNA Assembly* and *Assemble Multiple Fragments*. Through this action, the linearized vector becomes the first fragment in this virtual assembly.

6.10. Click on the Fragment dropdown menu and click on Number of Fragments. Change the number of fragments to 4 (vector + 3 U6-gRNA cassettes). Select the *Use directly as Fragment 1 option*.

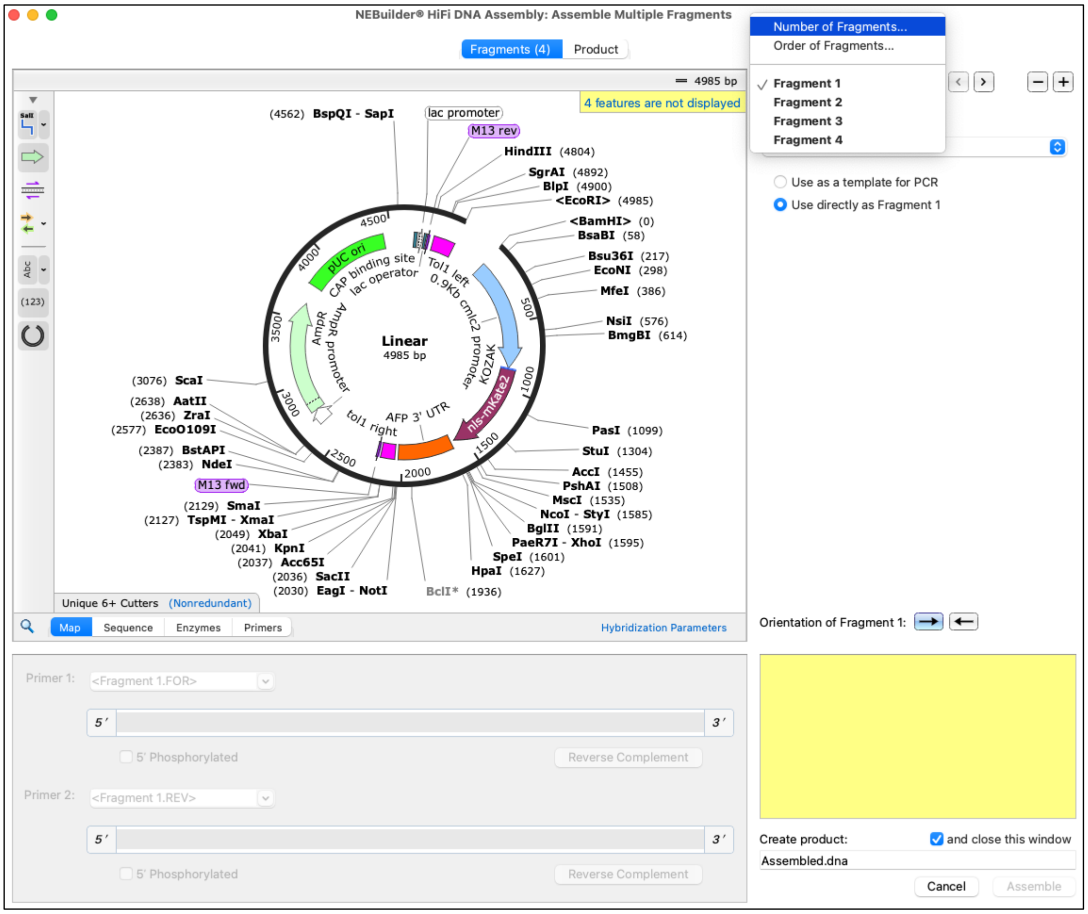

6.11. Click on the > button to move to Fragment 2. Select the file containing the amplified U6a-GOI-gA cassette in the *Source of Fragment 2* dropdown menu. Select the *Use directly as Fragment 2 option*.

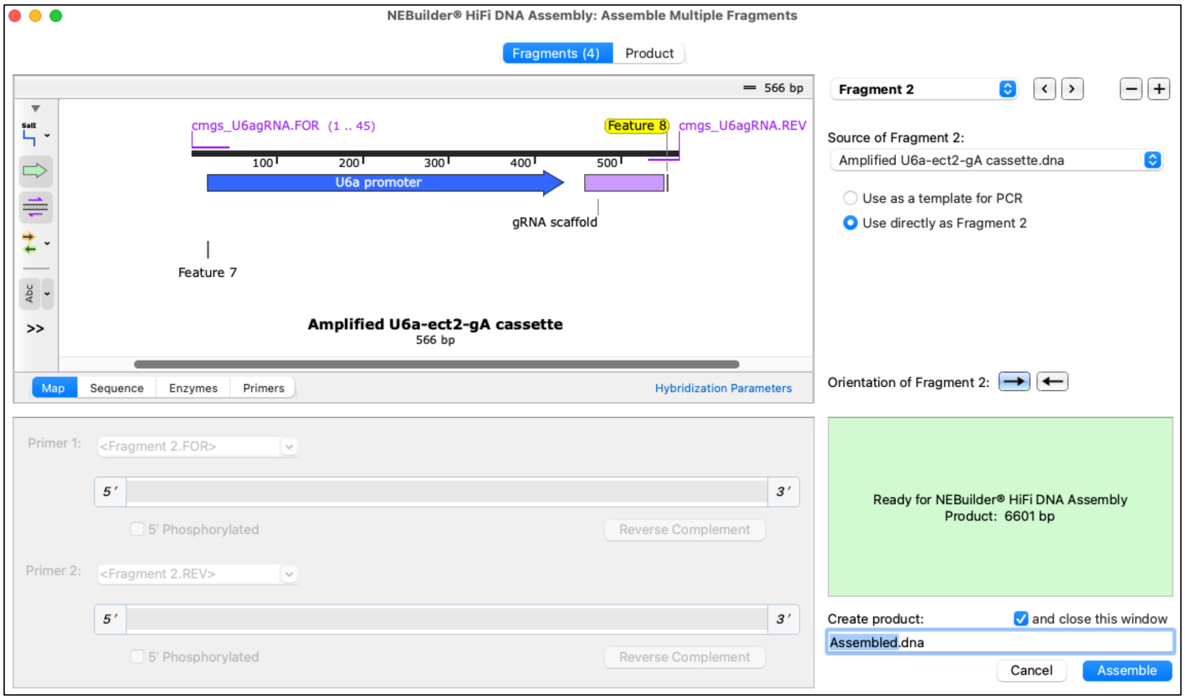

6.12. Following the same directions, add the U6b-GOI-gB cassette as Fragment 3 and U6c-GOI-gC as cassette 4.

6.13. Click on the *Product* Tab, type a new name for the final guide shuttle and click on *Assemble*.

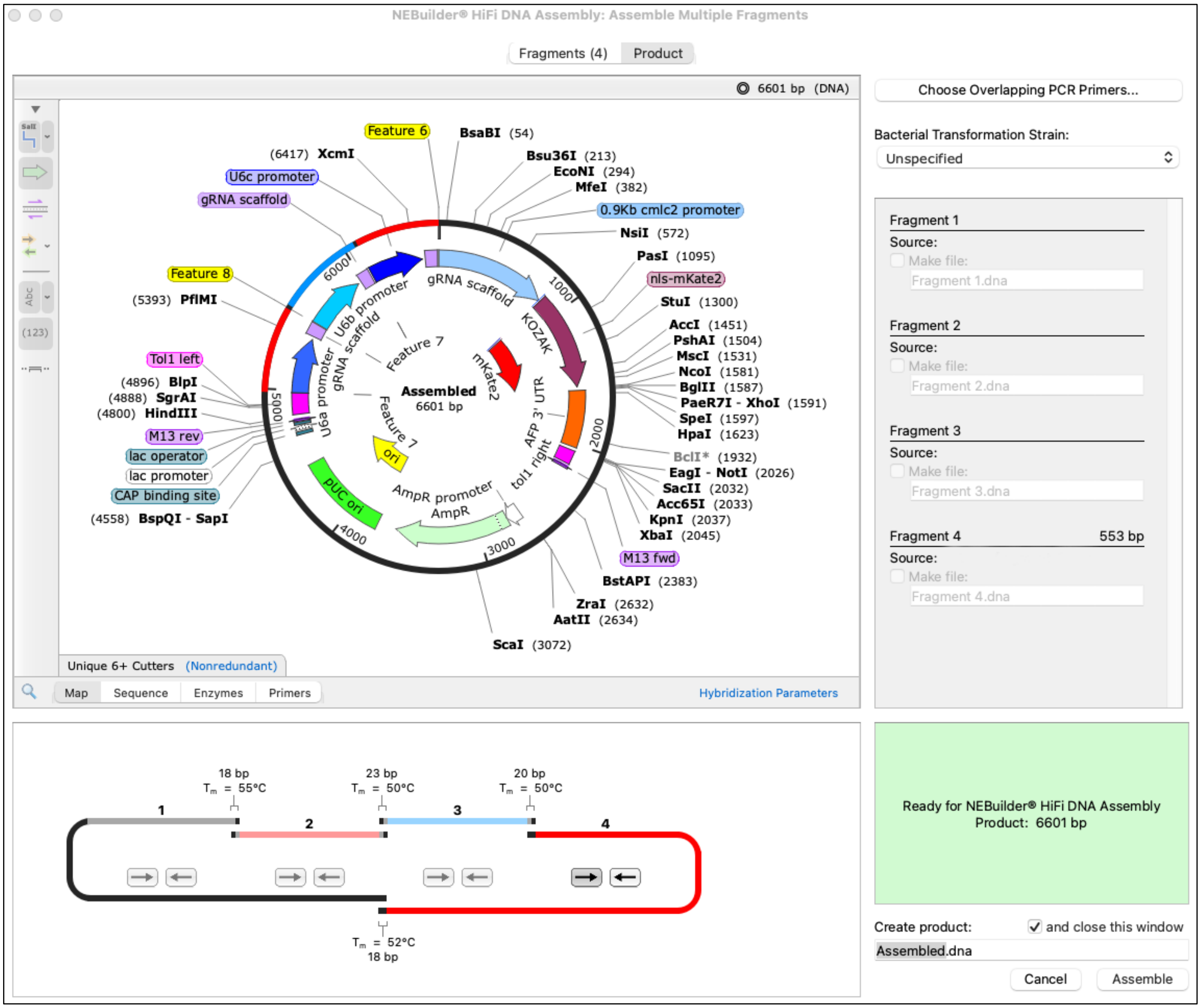


**<u>Method 2</u>:** If a Snapgene license is not available, the final sequence of the pU6-gRNA plasmids and final guide shuttles can be easily predicted by inserting the 18-nt target sequences (from step 1.6) in the following text files:

**<u>pU6a:sgRNA</u> (Addgene #64245, from PMID: 25855067)**

M13.FOR
**U6a promoter**
Target-specific sequence ***<u>replace by GG+your 18-nt target</u>* (from step 1.6)**
<u>gRNA scaffold</u>

```
ttgagatcctttttttctgcgcgtaatctgctgcttgcaaacaaaaaaaccaccgctaccagcggtggtttgtttgccggatcaagagc
taccaactctttttccgaaggtaactggcttcagcagagcgcagataccaaatactgtccttctagtgtagccgtagttaggccaccac
ttcaagaactctgtagcaccgcctacatacctcgctctgctaatcctgttaccagtggctgctgccagtggcgataagtcgtgtcttac
cgggttggactcaagacgatagttaccggataaggcgcagcggtcgggctgaacggggggttcgtgcacacagcccagcttggagcgaa
cgacctacaccgaactgagatacctacagcgtgagcattgagaaagcgccacgcttcccgaagggagaaaggcggacaggtatccggta
agcggcagggtcggaacaggagagcgcacgagggagcttccagggggaaacgcctggtatctttatagtcctgtcgggtttcgccacct
ctgacttgagcgtcgatttttgtgatgctcgtcaggggggcggagcctatggaaaaacgccagcaacgcggcctttttacggttcctgg
ccttttgctggccttttgctcacatgttctttcctgcgttatcccctgattctgtggataaccgtattaccgcctttgagtgagctgat
accgctcgccgcagccgaacgaccgagcgcagcgagtcagtgagcgaggaagcggaagagcgcccaatacgcaaaccgcctctccccgc
gcgttggccgattcattaatgcagctggcacgacaggtttcccgactggaaagcgggcagtgagcgcaacgcaattaatacgcgtaccg
ctagccaggaagagtttgtagaaacgcaaaaaggccatccgtcaggatggccttctgcttagtttgatgcctggcagtttatggcgggc
gtcctgcccgccaccctccgggccgttgcttcacaacgttcaaatccgctcccggcggatttgtcctactcaggagagcgttcaccgac
```

aaacaacagataaaacgaaaggcccagtcttccgactgagcctttcgttttatttgatgcctggcagttccctactctcgcgttaacgc
tagcatggatgttttcccagtcacgacgttgtaaaacgacggccagtcttaagctcgggccccggtctctcagtaagtctctcagcgttt
ctccagcctcggtcattcagtaagacgagtttcctgctctcggcgtcttttgttctggtcttcaaggagggggaaatgttctgcgcat
gcctgtgggggagggagaaggacacgtcactgaaaacgtccctgcatcacaccgagacacccaatcactcaagccaagaccagataatt
ttgcatatgctttacagtttgaaaattaccacggtaatccctcacacaaactctggatttgagatccttcaggttttatcagtttgcag
gtttatgtcaccatgatatagggtcagacttgatctaagggagctgaataagtggtttagtcactcaccacctcccaaaaacatacca
gaagtccctggtatatatagctctccctccagctcttggttcGGNNNNNNNNNNNNNNNNNNgtttaagagctatgctggaaacagcat
agcaagtttaaataaggctagtccgttatcaacttgaaaaagtggcaccgagtcggtgctttttttagtcagagacctctagaccagcca
ggacagaaatgcctcgacttcgctgctacccaaggttgccgggtgacgcacaccgtggaaacggatgaaggcacgaacccagtggacat
aagcctgttcggttcgtaagctgtaatgcaagtagcgtatgcgctcacgcaactggtccagaaccttgaccgaacgcagcggtggtaac
ggcgcagtggcggttttcatggcttgttatgactgtttttttggggtacagtctatgcctcgggcatccaagcagcaagcgcgttacgc
cgtgggtcgatgtttgatgttatggagcagcaacgatgttacgcagcagggcagtcgccctaaaacaaagttaaacattatgagggaag
cggtgatcgccgaagtatcgactcaactatcagaggtagttggcgtcatcgagcgccatctcgaaccgacgttgctggccgtacatttg
tacggctccgcagtggatggcggcctgaagccacacagtgatattgatttgctggttacggtgaccgtaaggcttgatgaaacaacgcg
gcgagctttgatcaacgaccttttggaaacttcggcttcccctggagagagcgagattctccgcgctgtagaagtcaccattgttgtgc
acgacgacatcattccgtggcgttatccagctaagcgcgaactgcaatttggagaatggcagcgcaatgacattcttgcaggtatcttc
gagccagccacgatcgacattgatctggctatcttgctgacaaaagcaagagaacatagcgttgccttggtaggtccagcggcggagga
actctttgatccggttcctgaacaggatctatttgaggcgctaaatgaaaccttaacgctatggaactcgccgcccgactgggctggcg
atgagcgaaatgtagtgcttacgttgtcccgcatttggtacagcgcagtaaccggcaaaatcgcgccgaaggatgtcgctgccgactgg
gcaatggagcgcctgccggcccagtatcagcccgtcatacttgaagctagacaggcttatcttggacaagaagaagatcgcttggcctc
gcgcgcagatcagttggaagaatttgtccactacgtgaaaggcgagatcaccaaggtagtcggcaaataaccctcgagccacccatgac
caaaatcccttaacgtgagttacgcgtcgttccactgagcgtcagaccccgtagaaaagatcaaaggatcttc

**<u>pU6b:sgRNA</u> (Addgene #64247, from PMID: 25855067)**

M13.FOR

**U6b promoter**

Target-specific sequence ***<u>replace by GG+your 18-nt target</u>* (from step 1.6)**

gRNA scaffold

ttgagatcctttttttctgcgcgtaatctgctgcttgcaaacaaaaaaaccaccgctaccagcggtggtttgtttgccggatcaagagc
taccaactctttttccgaaggtaactggcttcagcagagcgcagataccaaatactgtccttctagtgtagccgtagttaggccaccac
ttcaagaactctgtagcaccgcctacatacctcgctctgctaatcctgttaccagtggctgctgccagtggcgataagtcgtgtcttac
cgggttggactcaagacgatagttaccggataaggcgcagcggtcgggctgaacggggggttcgtgcacacagcccagcttggagcgaa
cgacctacaccgaactgagatacctacagcgtgagcattgagaaagcgccacgcttcccgaagggagaaaggcggacaggtatccggta
agcggcagggtcggaacaggagagcgcacgagggagcttccagggggaaacgcctggtatctttatagtcctgtcgggtttcgccacct
ctgacttgagcgtcgatttttgtgatgctcgtcaggggggcggagcctatggaaaaacgccagcaacgcggcctttttacggttcctgg
ccttttgctggccttttgctcacatgttctttcctgcgttatcccctgattctgtggataaccgtattaccgcctttgagtgagctgat
accgctcgccgcagccgaacgaccgagcgcagcgagtcagtgagcgaggaagcggaagagcgcccaatacgcaaaccgcctctccccgc
gcgttggccgattcattaatgcagctggcacgacaggtttcccgactggaaagcgggcagtgagcgcaacgcaattaatacgcgtaccg
ctagccaggaagagtttgtagaaacgcaaaaaggccatccgtcaggatggccttctgcttagtttgatgcctggcagtttatggcgggc
gtcctgcccgccaccctccgggccgttgcttcacaacgttcaaatccgctcccggcggatttgtcctactcaggagagcgttcaccgac
aaacaacagataaaacgaaaggcccagtcttccgactgagcctttcgttttatttgatgcctggcagttccctactctcgcgttaacgc
tagcatggatgttttcccagtcacgacgttgtaaaacgacggccagtcttaagctcgggccccggtctctctcattaccctccacgtgtc
tgtctgggttttcatgggctctgctctagtgagagcagcttccttttgtcggagtgttttgtgctcttatccaaggcaggggggatctg
cgcatgcctgtgggggggaggagaaggacacgtgaacaaaagctcctcgatgtcacacaggaagttcaggaactcatccaatcactctaa
agaaacggcctgtttccttcgcatacgcttacagctccaaaactctacggtaaacctacataaactgctggttttcaattttaaagaa
tttaagggtttacaggtttactactacacagtgatttactgacacatgtaagtgtaaatgagttgaataagtaagtaagccatatacca
cacatgaaacacatacccagaagtcactggtatatatagccgtcctccagactcccagttcGGNNNNNNNNNNNNNNNNNNgtttaaga
gctatgctggaaacagcatagcaagtttaaataaggctagtccgttatcaacttgaaaaagtggcaccgagtcggtgcttttttttaaac
caaagagacctctagaccagccaggacagaaatgcctcgacttcgctgctacccaaggttgccgggtgacgcacaccgtggaaacggat
gaaggcacgaacccagtggacataagcctgttcggttcgtaagctgtaatgcaagtagcgtatgcgctcacgcaactggtccagaacct
tgaccgaacgcagcggtggtaacggcgcagtggcggttttcatggcttgttatgactgtttttttggggtacagtctatgcctcgggca
tccaagcagcaagcgcgttacgccgtgggtcgatgtttgatgttatggagcagcaacgatgttacgcagcagggcagtcgccctaaaac
aaagttaaacattatgagggaagcggtgatcgccgaagtatcgactcaactatcagaggtagttggcgtcatcgagcgccatctcgaac
cgacgttgctggccgtacatttgtacggctccgcagtggatggcggcctgaagccacacagtgatattgatttgctggttacggtgacc
gtaaggcttgatgaaacaacgcggcgagctttgatcaacgaccttttggaaacttcggcttcccctggagagagcgagattctccgcgc
tgtagaagtcaccattgttgtgcacgacgacatcattccgtggcgttatccagctaagcgcgaactgcaatttggagaatggcagcgca
atgacattcttgcaggtatcttcgagccagccacgatcgacattgatctggctatcttgctgacaaaagcaagagaacatagcgttgcc
ttggtaggtccagcggcggaggaactctttgatccggttcctgaacaggatctatttgaggcgctaaatgaaaccttaacgctatggaa
ctcgccgcccgactgggctggcgatgagcgaaatgtagtgcttacgttgtcccgcatttggtacagcgcagtaaccggcaaaatcgcgc
cgaaggatgtcgctgccgactgggcaatggagcgcctgccggcccagtatcagcccgtcatacttgaagctagacaggcttatcttgga

caagaagaagatcgcttggcctcgcgcgcagatcagttggaagaatttgtccactacgtgaaaggcgagatcaccaaggtagtcggcaa
ataaccctcgagccacccatgaccaaaatcccttaacgtgagttacgcgtcgttccactgagcgtcagaccccgtagaaaagatcaaag
gatcttc

**pU6c:sgRNA (Addgene #64248, from PMID: 25855067)**

M13.FOR
**U6c promoter**
Target-specific sequence ***replace by GG+your 18-nt target* (from step 1.6)**
gRNA scaffold

ttgagatcctttttttctgcgcgtaatctgctgcttgcaaacaaaaaaaccaccgctaccagcggtggtttgtttgccggatcaagagc
taccaactctttttccgaaggtaactggcttcagcagagcgcagataccaaatactgtccttctagtgtagccgtagttaggccaccac
ttcaagaactctgtagcaccgcctacatacctcgctctgctaatcctgttaccagtggctgctgccagtggcgataagtcgtgtcttac
cgggttggactcaagacgatagttaccggataaggcgcagcggtcgggctgaacggggggttcgtgcacacagcccagcttggagcgaa
cgacctacaccgaactgagatacctacagcgtgagcattgagaaagcgccacgcttcccgaagggagaaaggcggacaggtatccggta
agcggcagggtcggaacaggagagcgcacgagggagcttccagggggaaacgcctggtatctttatagtcctgtcgggtttcgccacct
ctgacttgagcgtcgatttttgtgatgctcgtcaggggggcggagcctatggaaaaacgccagcaacgcggcctttttacggttcctgg
ccttttgctggccttttgctcacatgttctttcctgcgttatcccctgattctgtggataaccgtattaccgcctttgagtgagctgat
accgctcgccgcagccgaacgaccgagcgcagcgagtcagtgagcgaggaagcggaagagcgcccaatacgcaaaccgcctctccccgc
gcgttggccgattcattaatgcagctggcacgacaggtttcccgactggaaagcgggcagtgagcgcaacgcaattaatacgcgtaccg
ctagccaggaagagtttgtagaaacgcaaaaaggccatccgtcaggatggccttctgcttagtttgatgcctggcagtttatggcgggc
gtcctgcccgccaccctccgggccgttgcttcacaacgttcaaatccgctcccggcggatttgtcctactcaggagagcgttcaccgac
aaacaacagataaaacgaaaggcccagtcttccgactgagcctttcgttttatttgatgcctggcagttccctactctcgcgttaacgc
tagcatggatgttttcccagtcacgacgttgtaaaacgacggccagtcttaagctcgggcccggtctc**tccaatccgagagtctgtgaa**
**tgtttcgatgggccgtttgctggcgtctgagcatgttgtcctctggtctgcggcgggggaaggtccgcgcatgcctgtgggggaaggcg**
**aaggacacgaggacaaaagatcagttacgtcacagaagaactcatccaatcacacaagcctaaaaccagcttcctctgcatatgcttta**
**cagttcaaaaatactacggtaaacctctacaaaactgctggtttcaaactttgaaggttttaagcgtttgcaggtttgccccgaagagg**
**tttactgtcatgtttgaggtaaatgagttgaataagtaggtttatccacttaccacatggcagaaacatacccagaagtcccgggttta**
**tatagcagttctccaggctcctagttc**GGNNNNNNNNNNNNNNNNNNgtttaagagctatgctggaaacagcatagcaagtttaaataa
ggctagtccgttatcaacttgaaaaagtggcaccgagtcggtgctttttttaaacctgagagacctctagaccagccaggacagaaatg
cctcgacttcgctgctacccaaggttgccgggtgacgcacaccgtggaaacggatgaaggcacgaacccagtggacataagcctgttcg
gttcgtaagctgtaatgcaagtagcgtatgcgctcacgcaactggtccagaaccttgaccgaacgcagcggtggtaacggcgcagtggc
ggttttcatggcttgttatgactgtttttttggggtacagtctatgcctcgggcatccaagcagcaagcgcgttacgccgtgggtcgat
gtttgatgttatggagcagcaacgatgttacgcagcagggcagtcgccctaaaacaaagttaaacattatgagggaagcggtgatcgcc
gaagtatcgactcaactatcagaggtagttggcgtcatcgagcgccatctcgaaccgacgttgctggccgtacatttgtacggctccgc
agtggatggcggcctgaagccacacagtgatattgatttgctggttacggtgaccgtaaggcttgatgaaacaacgcggcgagctttga
tcaacgaccttttggaaacttcggcttcccctggagagagcgagattctccgcgctgtagaagtcaccattgttgtgcacgacgacatc
attccgtggcgttatccagctaagcgcgaactgcaatttggagaatggcagcgcaatgacattcttgcaggtatcttcgagccagccac
gatcgacattgatctggctatcttgctgacaaaagcaagagaacatagcgttgccttggtaggtccagcggcggaggaactctttgatc
cggttcctgaacaggatctatttgaggcgctaaatgaaaccttaacgctatggaactcgccgcccgactgggctggcgatgagcgaaat
gtagtgcttacgttgtcccgcatttggtacagcgcagtaaccggcaaaatcgcgccgaaggatgtcgctgccgactgggcaatggagcg
cctgccggcccagtatcagcccgtcatacttgaagctagacaggcttatcttggacaagaagaagatcgcttggcctcgcgcgcagatc
agttggaagaatttgtccactacgtgaaaggcgagatcaccaaggtagtcggcaaataaccctcgagccacccatgaccaaaatccctt
aacgtgagttacgcgtcgttccactgagcgtcagaccccgtagaaaagatcaaaggatcttc

**pTol1 cm:n-mKate guide shuttle**

Tol1 arms
**U6a promoter**
**U6b promoter**
U6c promoter
target-specific sequence ***replace by GG+your 18-nt targets* (from step 1.6)**
gRNA scaffold
**cmlc2 promoter** + n-mKate2 (reporter)

gtagaaaagatcaaaggatcttcttgagatcctttttttctgcgcgtaatctgctgcttgcaaacaaaaaaaccaccgctaccagcggt
ggtttgtttgccggatcaagagctaccaactctttttccgaaggtaactggcttcagcagagcgcagataccaaatactgttcttctag
tgtagccgtagttaggccaccacttcaagaactctgtagcaccgcctacatacctcgctctgctaatcctgttaccagtggctgctgcc
agtggcgataagtcgtgtcttaccgggttggactcaagacgatagttaccggataaggcgcagcggtcgggctgaacggggggttcgtg
cacacagcccagcttggagcgaacgacctacaccgaactgagatacctacagcgtgagctatgagaaagcgccacgcttcccgaaggga
gaaaggcggacaggtatccggtaagcggcagggtcggaacaggagagcgcacgagggagcttccagggggaaacgcctggtatctttat
agtcctgtcgggtttcgccacctctgacttgagcgtcgatttttgtgatgctcgtcaggggggcggagcctatggaaaaacgccagcaa

cgcggcctttttacggttcctggccttttgctggccttttgctcacatgttctttcctgcgttatcccctgattctgtggataaccgta
ttaccgcctttgagtgagctgataccgctcgccgcagccgaacgaccgagcgcagcgagtcagtgagcgaggaagcggaagagcgccca
atacgcaaaccgcctctccccgcgcgttggccgattcattaatgcagctggcacgacaggtttcccgactggaaagcgggcagtgagcg
caacgcaattaatgtgagttagctcactcattaggcaccccaggctttacactttatgcttccggctcgtatgttgtgtggaattgtga
gcggataacaatttcacacaggaaacagctatgaccatgattacgccaagcttgcatggtccctttagccagtagcggttctaggcacg
ggccgtccgggcggtggcctggggcggaaaactgaagggggcggcaccggcggctcagcccttgtaatatattatatgcaccactatt
ggtttacttatgtcacagtttgtaagtttgtaacagcctgaacctggatcgtcagtaagtctctcagcgtttctccagcctcggtcatt
cagtaagacgagtttcctgctctcggcgtcttttgttctggtcttcaaggagggggaaatgttctgcgcatgcctgtgggggagggag
aaggacacgtcactgaaaacgtccctgcatcaccgagacacccaatcactcaagccaagaccagataattttgcatatgctttacag
tttgaaaattaccacggtaatccctcacacaaactctggatttgagatccttcaggttttatcagtttgcaggtttatgtcaccatgat
atagggtcagacttgatctaagggagctgaataagtggtttagtcactcaccacctcccaaaaacatacccagaagtccctggtatata
tagctctccctccagctcttggttcGGNNNNNNNNNNNNNNNNNNgtttaagagctatgctggaaacagcatagcaagtttaaataagg
ctagtccgttatcaacttgaaaaagtggcaccgagtcggtgctttttttagtcatctcattaccctccacgtgtctgtctgggttttcat
gggctctgctctagtgagagcagcttccttttgtcggagtgttttgtgctcttatccaaggcaggggggatctgcgcatgcctgtgggg
ggaggagaaggacacgtgaacaaaagctcctcgatgtcacacaggaagttcaggaactcatccaatcactctaaagaaacggcctgttt
ccttcgcatacgcttacagctccaaaactctacggtaaacctacataaactgctggttttcaattttaaagaatttaagggtttacag
gtttactactacacagtgatttactgacacatgtaagtgtaaatgagttgaataagtaagtaagccatataccacacatgaaacacata
cccagaagtcactggtatatatagccgtcctccagactcccagttcGGNNNNNNNNNNNNNNNNNNgtttaagagctatgctggaaaca
gcatagcaagtttaaataaggctagtccgttatcaacttgaaaaagtggcaccgagtcggtgctttttttaaaccaaatccaatccgag
agtctgtgaatgtttcgatgggccgtttgctggcgtctgagcatgttgtcctctggtctgcggcgggggaaggtccgcgcatgcctgtg
ggggaaggcgaaggacacgaggacaaaagatcagttacgtcacagaagaactcatccaatcacacaagcctaaaaccagcttcctctgc
atatgctttacagttcaaaaatactacggtaaacctctacaaaactgctggtttcaaactttgaaggttttaagcgtttgcaggtttgc
cccgaagaggtttactgtcatgtttgaggtaaatgagttgaataagtaggtttatccacttaccacatggcagaaacatacccagaagt
cccgggtttatatagcagttctccaggctcctagttcGGNNNNNNNNNNNNNNNNNNgtttaagagctatgctggaaacagcatagcaa
gtttaaataaggctagtccgttatcaacttgaaaaagtggcaccgagtcggtgctttttttaaacctgacctccaaatcagcagactta
acattctttaaaatgattgattcaatagtgataaaaatcaggcatagccagttgtaactttagataaattacagaaaatgtcaaataca
gagaaccgattctttttttatgatacatccaagcacacatttaacacaatccaggcaaaccccgaatttcacagtcacaagcactgtttg
tacaagagctttgcctaaggacacacagtctctataagtccaggtcgttggtttcactcttattttaaacatgtgacatttttcctgcc
atcctgtcttaggctgctgtttgcttcattccatgtcacattaatttcctcagtagcaccttttacacacacagccaatcttttccaga
aaattcaattgctttgaaaagataatgtgtgaacaaatccatttagaaaaggaaaattaagaatttgtaaaatcatctgtaaattgttg
gcattcttctgtatatgaacatcacatcatttacaggtaaaggtctggtcattaattatatgacaatttactggtattattttgtgaaa
ggggctattttcaatgcattcatccatccttttcatccctcaaatctctcattcacgtcccccctccccatctgcacactttatctcatt
ttccaccctgctggaatctgagcacttgtgcagttatcagggctcctgtatttaggaggctctgggtgtccatgtaggggacgaacaga
aacactgcagacctttatagaagaacaaatgataagagtcctcatacataaagactccattagaaacgtcagtgacccaggagcccaga
ccaacagcaaagcagacagtgaaccgccaccatgccaaagaagaagcgtaaggtagtgagcgagctgattaaggagaacatgcacatga
agctgtacatggagggcaccgtgaacaaccaccacttcaagtgcacatccgagggcgaaggcaagccctacgagggcacccagaccatg
agaatcaaggcggtcgagggcggccctctccccttcgccttcgacatcctggctaccagcttcatgtacggcagcaaaaccttcatcaa
ccacacccagggcatccccgacttctttaagcagtccttccccgagggcttcacatgggagagagtcaccacatacgaagacgggggcg
tgctgaccgctacccaggacaccagcctccaggacggctgcctcatctacaacgtcaagatcagaggggtgaacttcccatccaacggc
cctgtgatgcagaagaaaacactcggctgggaggcctccaccgagaccctgtaccccgctgacggcggcctggaaggcagagccgacat
ggccctgaagctcgtgggcgggggccacctgatctgcaacttgaagaccacatacagatccaagaaacccgctaagaacctcaagatgc
ccggcgtctactatgtggacagaagactggaaagaatcaaggaggccgacaaagagacctacgtcgagcagcacgaggtggctgtggcc
agatactgcgacctccctagcaaactggggcacagagtttaaagccaccatggagatctcgagactagtaattaagtctcagccaccgt
taactgaacatgtcaaaacctgtggagactgttgagatttgatgttctgaaaagataaagcctataaataaaatgttgcccaaatttcc
tgcctgatgtttttctttgtctttgctacatggctttgctgctcggatcggctcactctgtgtatgccacgttcactttgtactctcct
tctcacggtaggtttattattttagatgtgcagttagtttctgtgaaataacacaccacacactgatattgtctgtgcattgacttgg
tgagtgcacattgtttttgatcttgacatatttatatttgattgatcaggtgaactgtgtgaatctaaagtgctccatacagatgttct
gcattgaaaatattctcattttattagtggaagtgagtgtatgccagcggccgcggtaccctgcagtctagacatttcagttgacgaag
acaaacaaagttctgttgtgactatgggggggggggcgcctggggatggtctcgcccggggagtaattcaggtagaaccgccactgc
ctttagcgacattcactggccgtcgttttacaacgtcgtgactgggaaaaccctggcgttacccaacttaatcgccttgcagcacatcc
ccctttcgccagctggcgtaatagcgaagaggcccgcaccgatcgcccttcccaacagttgcgcagcctgaatggcgaatggcgcctga
tgcggtattttctccttacgcatctgtgcggtatttcacaccgcatatggtgcactctcagtacaatctgctctgatgccgcatagtta
agccagccccgacacccgccaacacccgctgacgcgccctgacgggcttgtctgctcccggcatccgcttacagacaagctgtgaccgt
ctccgggagctgcatgtgtcagaggttttcaccgtcatcaccgaaacgcgcgagacgaaagggcctcgtgatacgcctatttttatagg
ttaatgtcatgataataatggtttcttagacgtcaggtggcacttttcggggaaatgtgcgcggaacccctatttgtttatttttctaa
atacattcaaatatgtatccgctcatgagacaataaccctgataaatgcttcaataatattgaaaaaggaagagtatgagtattcaaca
tttccgtgtcgcccttattcccttttttgcggcattttgccttcctgtttttgctcacccagaaacgctggtgaaagtaaaagatgctg
aagatcagttgggtgcacgagtgggttacatcgaactggatctcaacagcggtaagatccttgagagttttcgccccgaagaacgtttt
ccaatgatgagcacttttaaagttctgctatgtggcgcggtattatcccgtattgacgccgggcaagagcaactcggtcgccgcataca
ctattctcagaatgacttggttgagtactcaccagtcacagaaaagcatcttacggatggcatgacagtaagagaattatgcagtgctg
ccataaccatgagtgataacactgcggccaacttacttctgacaacgatcggaggaccgaaggagctaaccgcttttttgcacaacatg
ggggatcatgtaactcgccttgatcgttgggaaccggagctgaatgaagccataccaaacgacgagcgtgacaccacgatgcctgtagc

aatggcaacaacgttgcgcaaactattaactggcgaactacttactctagcttcccggcaacaattaatagactggatggaggcggata
aagttgcaggaccacttctgcgctcggcccttccggctggctggtttattgctgataaatctggagccggtgagcgtgggtctcgcggt
atcattgcagcactggggccagatggtaagccctcccgtatcgtagttatctacacgacggggagtcaggcaactatggatgaacgaaa
tagacagatcgctgagataggtgcctcactgattaagcattggtaactgtcagaccaagtttactcatatatactttagattgatttaa
aacttcatttttaatttaaaaggatctaggtgaagatcctttttgataatctcatgaccaaaatcccttaacgtgagttttcgttccac
tgagcgtcagacccc
